# Tracing the invasion of a leaf-mining moth in the Palearctic through DNA barcoding of historical herbaria

**DOI:** 10.1101/2021.10.07.463492

**Authors:** Natalia I. Kirichenko, Evgeny V. Zakharov, Carlos Lopez-Vaamonde

## Abstract

Historical herbaria are valuable sources of data in invasion biology. Here we study the invasion history of the lime leaf-miner, *Phyllonorycter issikii*, by surveying over 15 thousand herbarium specimens of limes (*Tilia* spp.) collected in the Palearctic during last 253 years (1764–2016). The majority of herbarium specimens with the pest’s mines (89%) originated from East Asia (1859–2015), whereas remaining 11% of specimens with the mines came from Europe, European Russia and Western Siberia (1987–2015). These results support the hypothesis of a recent *Ph. issikii* invasion from Eastern to Western Palearctic.

Single molecule real-time sequencing of the COI barcode region of 93 archival larvae and pupae (7–162 years old) dissected from the mines on historical herbaria allowed to distinguish between *Ph. issikii* and *Ph. messaniella*, a polyphagous species rarely feeding on *Tilia*, which mines were found in herbarium from Europe dated by 1915–1942. We discovered 25 haplotypes of *Ph. issikii*, of which 16 haplotypes were present solely in East Asia, and revealed wide distribution of the species in China. Six haplotypes shared between Eastern and Western Palearctic suggest the contribution of *Ph. issikii* populations from the Russian Far East, China and Japan to the westward invasion.

## Introduction

Herbarium specimens collected over the centuries in different biogeographic regions have a great value for science^1,2^. Indeed, they represent not only an important source of data for botanists, but they also provide entomologists with paramount information on past diversity, distribution, abundance, and trophic associations of insects^3,4,5,6^.

The role of herbaria in studying the distribution of alien phytophagous insects has been underestimated for long time^7,8,9^. Despite the fact that botanists tend to avoid collecting damaged plant parts to herbaria^10^, the traces of endophagous insects, in particular leaf miners (i.e. cavities of different shapes in leaves that do not visually disturb the integrity of the leaf lamina) are often present in pressed plant specimens^11^. Moreover, some of those leaf mines in pressed leaves containing larvae and pupae of leaf-mining insects have been used for various studies including systematics^12^, phylogeography^4^, island biogeography and conservation^3,7,13^. Molecular analyses of archival samples helped to reveal the area of origin of invasive insects, such as the horse-chestnut leaf miner, *Cameraria ohridella* Deschka and Dimić, 1986 (Lepidoptera: Gracillariidae), confirming its Balkan origin^4^.

The leaf mining moth, *Phyllonorycter issikii* (Kumata, 1963) (Lepidoptera: Gracillariidae) is the only species of this genus that feeds exclusively on limes, *Tilia* spp. (Malvaceae) in the Palearctic. In the last few decades, this species, which was originally known from East Asia (Japan, Korea, Russian Far East)^14,15,16,17^, invaded Western Palearctic and became a pest of lime trees^18,19,20,21,22^. In a recent analysis of COI mtDNA barcodes of 377 *P. issikii*’s specimens collected across the modern rage in the Palearctic, we revealed unusually high genetic diversity of the species in the recently invaded region (Western Palearctic) compared to the putative native region (Eastern Palearctic, in particular Eastern Asia) questioning the hypotheses about the moth’s region of origin and geographic expansion^20^.

To test the hypothesis that *Ph. issikii* is native to East Asia and has recently invaded Western Palearctic, we examined pressed lime (*Tilia*) specimens collected across the Palearctic in the last 253 years. We expected to find archival leaf mines in herbaria specimens in Eastern Asia well before the beginning of the invasion and in Western Europe only after the invasion event. We also took advantage of the recent advances in DNA barcoding of samples with degraded DNA^22,23^ to analyse larvae and pupae found inside archival leaf mines to understand the genetics of invasion and explain the high level of haplotype diversity in the neocolonized area in Europe.

Additionally, we surveyed herbaria from the Nearctic dated by 1850–2010 and DNA barcoded archival larvae and pupae (10–170 years old) to test whether *Ph. issikii* occurs in that biogeographic region.

## Results

### Detection of *Phyllonorycter* mines in historical herbaria

Only 1.5% (226 out of 15,009) of herbarium specimens of *Tilia spp*. examined from the Palearctic contained characteristic *Ph. issikii* leaf mines. These records corresponded to 185 geographical locations across the Palearctic, with the most western point in Germany (Hessen; the herbarium specimen dated by 2004) to the most eastern locations in Japan (on the island Hokkaido; 1885–1974) (Fig. 1).

**Figure 1.**
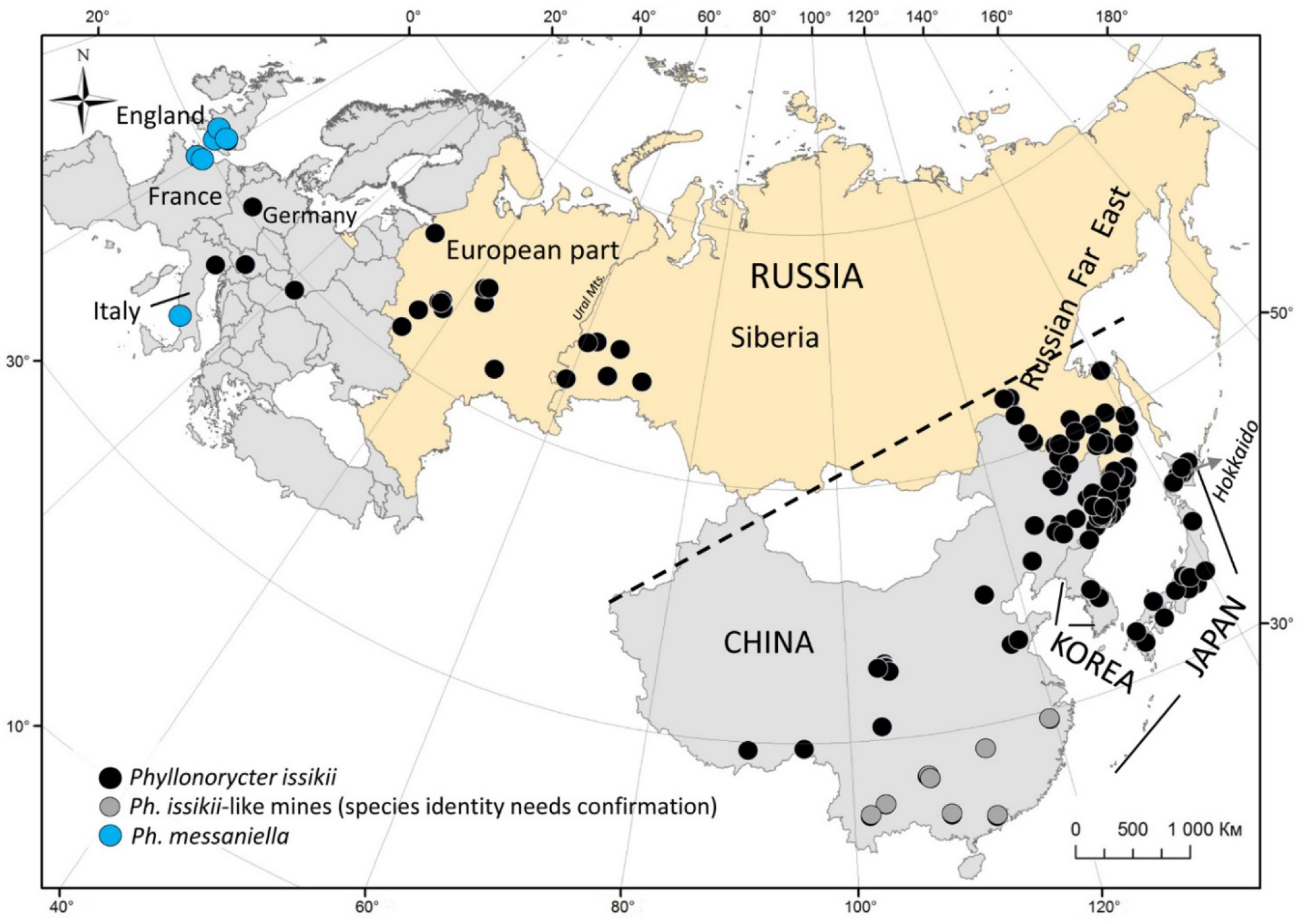
The localities where herbarium specimens of *Tilia* spp. carrying typical *Phyllonorycter* mines were collected in the Palearctic in the last 253 years. The dotted line presumably divides *Ph. issikii* range to native (below the line) and invaded (above the line).

Most specimens with leaf mines (90%; 204/226) and most leaf mines (89.6%; 1,154/1,288) originated from the primary range in East Asia, in particular from the Russian Far East (RFE) (Fig. 2a). In some cases, leaves were significantly attacked, with up to 12 mines per leaf. On the other hand, we found only 22 (10%; 22/226) specimens with 134 mines (10%; 134/1,288) from the invaded region in Western Palearctic, with the majority from European Russia (Fig. 2b).

**Figures 2.**
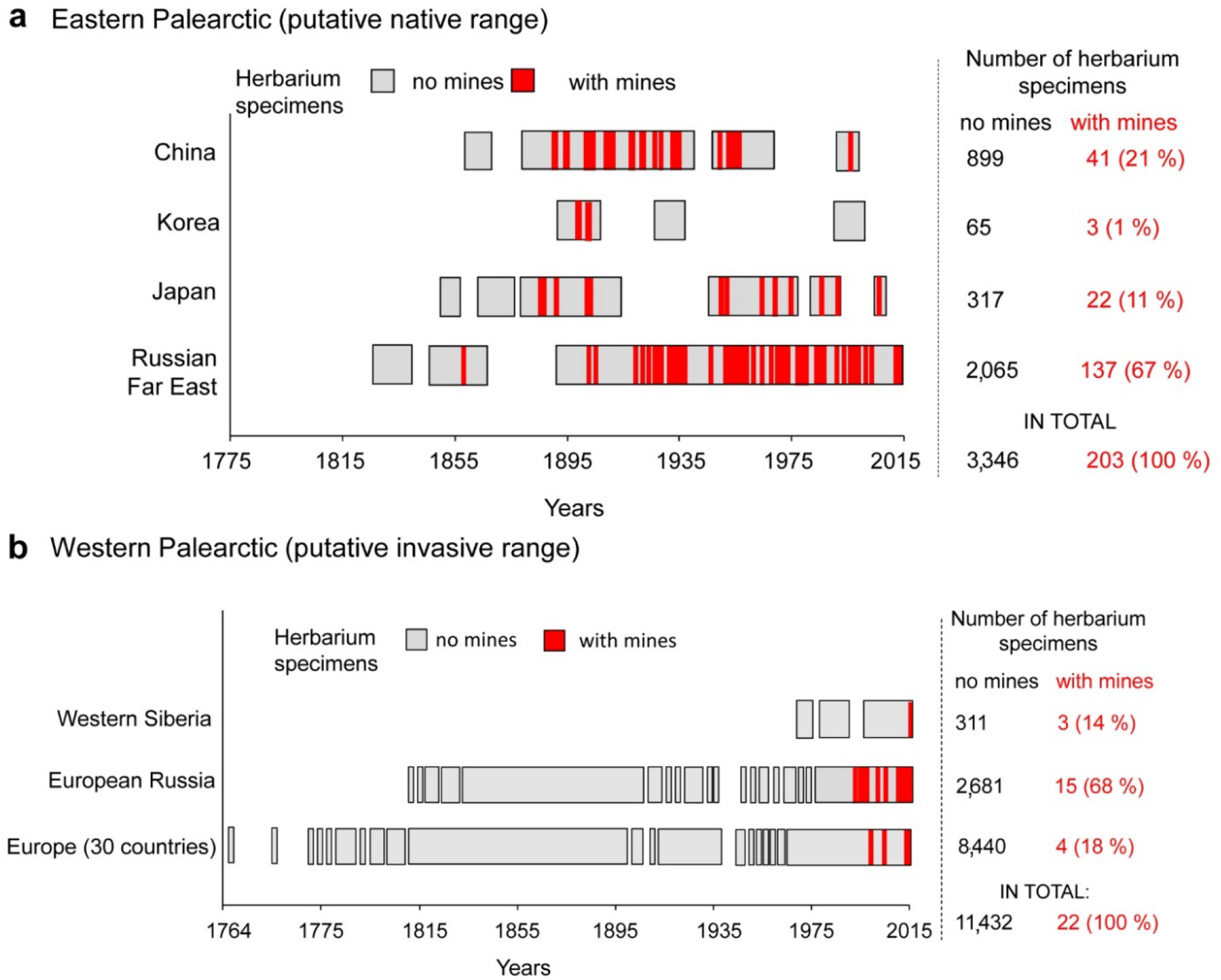
The presence of the typical *Phyllonorycter issikii* mines in the herbarium specimens collected in the putative native range (**a**) and putative invasive range (**b**) over the past 253 years. The number of herbarium specimens with and without mines and the percentage of the specimens with mines from the total number of specimens with mines (in brackets) are given next to each graph.

The average number of leaf mines per herbarium specimen found in native (5.65±0.77) and invaded regions (6.09±1.70) was not significantly different (Mann–Whitney U-test: U = 20145; Z = 0.43; *P* = 0.43). However, the infestation rate by *Ph. issikii*, i.e. percentage of leaves with mines per herbarium specimen was statistically higher in the West than in the East: 35% ± 8.19 vs. 23% ± 1.94 (Mann–Whitney U-test: U = 1339; Z = 2.30; *P* = 0.02). Leaf mines from the East were significantly older than those from the West (Mann–Whitney U-test: U = 81; Z = –4.4; *P* < 0.001). Indeed, *Ph. issikii* mines from the West were found on herbarium specimens collected exclusively in the last three decades (1987–2015), whereas in East Asia, they were detected in herbarium samples dating back to 1859 (Fig. S1). The oldest mines were revealed in the pressed lime leaves sampled in the RFE in Amur Oblast in 1859, i.e. 162 years ago and 104 years before the date of *Ph. issikii* description from Japan (the year 1963). The time lag between the earliest mine record on the East (1859) and that on the West (1987) is 128 years.

We found six European herbarium specimens with eleven mines of another species, polyphagous moth *Ph. messaniella* (Fig. 1; Supplementary Table S1). Seven out of eleven mines were assigned to *Ph. messaniella* by their morphology, i.e. presence of one distinct longitude fold on the epidermis covering the mines. However, the remaining four mines were at an early stage of development and the fold was not visible therefore DNA barcoding was used to identify them (see results below).

None of the 683 North American herbarium specimens of *Tilia* contained *Ph. issikii* mines. However, we found 37 specimens (5.4%) with 88 leaf mines of *Ph. lucetiella* and *Ph. tiliacella* with the specimens’ ages ranging from 11 to 197 years old (Supplementary Table S2).

### Distribution of *Ph. issikii* in the past

#### Eastern Palearctic (native range)

We found *Ph. issikii* mines on 203 herbarized lime specimens in the Eastern Palearctic: on 137 herbarium specimens from the RFE, 41 from China, 22 from Japan, and 3 from Korea (Fig. 2a).

In Japan, from where *Ph. issikii* was formally described^14^, the mines were found in herbaria samples collected between 1886 and 2011, from the North to the South (32–45°N) and from the West to the East (130–145°E) (Fig. 3a). The earliest finding of *Ph. issikii* mines dated back to 1886 from Hokkaido.

**Figure 3.**
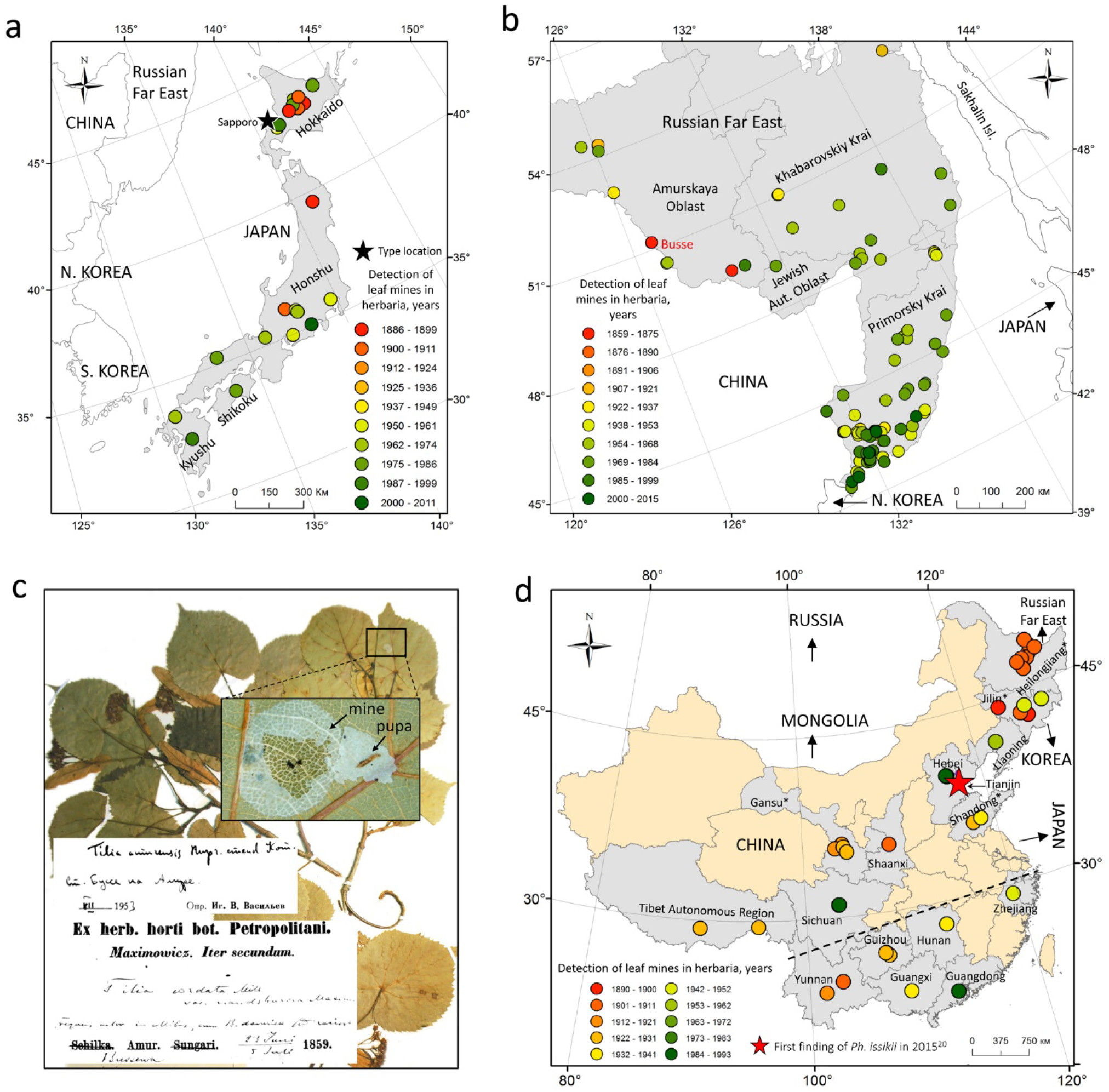
Past distribution of *Phyllonorycter issikii* in Japan (**a**), the Russian Far East (**b**), and China (**d**) based on findings of leaf mines in herbaria collected in 1859–2015. The village Busse is the location of the earliest finding of *Ph. issikii* mines (1859) (**b, c**). In China (**d**), the provinces where typical mines were found in herbaria are shaded in gray; in the provinces marked by an asterisk, the identification of *Ph. issikii* archival specimens was confirmed by DNA barcoding. The dotted line shows a schematical border between the Palearctic (above the line) and the Indomalaya (below the line).

In the RFE, herbarium samples with leaf mines were collected between 1859 and 2005 in 137 localities in Primorsky Krai, Khabarovsk Krai, Jewish Autonomous Oblast and Amur Oblast (Fig. 3b). The typical mines were also found in pressed lime specimens from the islands Russky, Popov and Askold (sampling years 1973–1998). According to the findings of mines in herbaria, the historical range of *Ph. issikii* in the RFE covered a significant area reaching the latitude of 42°–54.5° N and the longitude of 126°–139° E. Importantly, in the RFE the mines of *Ph. issikii* were regularly found in the herbaria from the localities next to the border with China and North Korea (Fig. 3b). The oldest herbarium specimen with *Ph. issikii* mines and with the species identity confirmed genetically (see DNA barcoding section below) was collected in 1859 (in village Busse, Amur Oblast, 1.5 km from the border with the Chinese province Heilongjiang) (Fig. 3b,c), which was the earliest confirmed finding of the moth signs in East Asia. Furthermore, the time lag between this earliest finding in herbaria (1859) and that of *Ph. issikii* specimens from the type series sampled in Japan in 1954^14^ was 95 years.

In China, the typical mines on pressed lime specimens were found in 15 provinces: from Heilongjiang in the northeast to Yunnan, Guangdong and Guangxi Zhuang Autonomous Region in the south, i.e. on the territory between 47–32° N and 105–125° E (Fig. 3d). The age of the herbarium specimens on which those mines were detected in China varied from 28 to 131 years (the herbarium specimens were collected in 1890–1993) (Fig. 3d). The oldest specimens with the moth mines originated from the northeastern provinces Jilin (1896) and Heilongjiang (1902).

In Korea, only six mines were found in three lime specimens: one mine in two herbarium specimens collected in 1900–1902 during the Korean-Sakhalin expedition by the Imperial Russian Geographical Society and the other four mines on the specimen dated by 1909 without indication of the exact sampling location on the label.

#### Western Palearctic (neocolonized range)

In Europe, no *Ph. issikii* mines were found in any of the 11,121 pressed lime specimens collected before 1987. Leaf mines of *Ph. issikii* were detected only in four pressed lime specimens, all originating from European Russia, with the earliest record from 1987 near the city Zlatoust in Chelyabinsk Oblast (the region on the border of Europe with Asia). The other three records corresponded to the Volga Region, in particular Samara Oblast (one herbarium specimen sampled in 1990), and the Urals, in particular Sverdlovsk Oblast (two herbarium specimens, 1991) (Table 1). All other records corresponded to the period between 2000 and 2016 (Table 1). In European Russia, the mines were found in herbaria specimens from eight distantly located regions (Table 1). Further west, in Europe, the mines were detected in herbarium specimens collected in Austria, Germany, Italy, and Slovakia (2006–2015) (Table 1). Further east, in Siberia, the mines were recorded exceptionally on pressed lime specimens originating from the western regions: Khanty-Mansi Autonomous Okrug and Tyumen Oblast (Table 1).

**Table 1.** Detected *Phyllonorycter issikii* mines in the lime herbarium specimens collected in 1987–2016 in different locations in Western Palearctic.

| № | Country, region<br>(Obl. – Oblast) | Location (n/d – no<br>data, vil. – village)<br>and biotope [f –<br>forest, p – park] | Host<br>plant* | Number of |  |  | Year |
| --- | --- | --- | --- | --- | --- | --- | --- |
|  |  |  |  | mines | leaves with<br>mines<br>[percentage<br>of mined<br>leaves] | leaves in<br>herbarium<br>specimen |  |
| EUROPE |  |  |  |  |  |  |  |
| 1 | Austria, Styrian | n/d; [f] | *T.c.* | 10 | 6 [11] | 54 | 2015 |
| 2 | Germany, Hessen | n/d; [f] | *T.c.* | 2 | 2 [9] | 22 | 2014 |
| 3 | Italy, Veneto | n/d; [f] | *T.p.* | 7 | 5 [45] | 11 | 2011 |
| 4 | Slovakia | vil. Trebejov; [f] | *T.c.* | 2 | 2 [11] | 18 | 2006 |
| RUSSIA: European part |  |  |  |  |  |  |  |
| 5 | Chelyabinsk Obl. | Zlatoust; [f] | *T.c.* | 3 | 3 [30] | 10 | 1987 |
| 6 | Kaluga Obl. | vil. Klykovo; [p] | *T.p.* | 1 | 1 [11] | 9 | 2008 |
| 7 | Kostroma Obl. | vil. Kozlovo; [f] | *T.c.* | 12 | 7 [28] | 25 | 2014 |
| 8 | Kostroma Obl. | vil. Bogorodskoe;<br>[f] | *T.c.* | 28 | 18 [100] | 18 | 2014 |
| 9 | Kostroma Obl. | vil. Seraphikha; [f] | *T.c.* | 5 | 5 [26] | 19 | 2013 |
| 10 | Kursk Obl. | Zheleznogorsk; [p] | *T.c.* | 26 | 6 [100] | 6 | 2011 |
| 11 | Leningrad Obl. | Pavlovsk; [p] | *T.p.* | 1 | 1 [11] | 9 | 2000 |
| 12 | Moscow Obl. | Moscow, Lublino;<br>[p] | *T.c.* | 2 | 2 [14] | 14 | 2012 |
| 13 | Moscow Obl. | vil. Gorodishe | *T.p.* | 3 | 2 [100] | 2 | 2015 |
| 14 | Moscow Obl. | Moscow [p] | *T.p.* | 2 | 2 [18] | 11 | 2012 |
| 15 | Moscow Obl. | Moscow [p] | *T.p.* | 17 | 8 [100] | 8 | 2006 |
| 16 | Moscow Obl. | Moscow [p] | *T.p.* | 1 | 1 [14] | 7 | 2006 |
| 17 | Samara Obl. | Zhiguli Nat.<br>Reserve; [f] | *T.c.* | 7 | 6 [33] | 18 | 1990 |
| 18 | Sverdlovsk Obl. | Sverdlovsk suburb<br>[f] | *T.c.* | 1 | 1 [8] | 13 | 1991 |
| 19 | Sverdlovsk Obl. | Sverdlovsk suburb<br>[f] | *T.c.* | 1 | 1 [4] | 26 | 1991 |
| RUSSIA: Western Siberia |  |  |  |  |  |  |  |
| 20 | Khanty-Mansi AO | vil. Kumisniy, [f] | *T.c.* | 1 | 1 [4] | 23 | 2015 |
| 21 | Tyumen Obl. | Vikulovskiy<br>district, [f] | *T.c.* | 1 | 1 [7] | 15 | 2007 |
| 22 | Tyumen Obl. | Isetskiy district, [f] | *T.c.* | 1 | 1 [5] | 19 | 2012 |

\**T.c.* – *Tilia cordata*, *T.p.* – *Tilia platyphillos*.

### Trophic associations of *Ph. issikii* with the limes in the past

We found mines of *Ph. issikii* in pressed specimens from 20 different *Tilia* species across the Palearctic: on 18 lime species in East and on two species in the West (Supplementary Fig. S2, Table S3).

In the East, 33% of all herbarium specimens with the mines were found on *T. amurensis* (74/226) followed by *T. taquetii* (19%, i.e. 42/226) and *T. mandshurica* (11%, 24/226) (Supplementary Fig. S2, Table S3). Occasionally, the mines were detected on other 15 different East Asian lime species (Supplementary Fig. S2, Table S3). We documented *Ph. issikii* mines on five novel host plants in China: *T. chinensis, T. intonsa, T. leptocarya, T. miqueliana*, and *T. paucicostata* (Supplementary Fig. S2). Among these five species, *T. leptocarya* is presently considered as a synonym of *T. endochrysea*. In the West, *Ph. issikii* mines were found in the herbarium specimens sampled on *T. cordata* and *T. platyphyllos*. In European Russia and Western Siberia, they were revealed only on *T. cordata* (Supplementary Fig. S2).

### DNA barcoding of archival samples

DNA barcodes were obtained for 88 out of 93 (i.e. 95%) archival larvae and pupae dissected from 7 up to 162-year-old pressed lime leaf mine specimens. The remaining five archival larvae and pupae (four 74–125-year-old specimens from the Palearctic, and one 171-year-old specimen from the Nearctic) failed to produce sequences. Among 88 sequenced specimens 73 specimens originated from the Palearctic (dated by 1859–2014) and 15 specimens were from the Nearctic (dated by 1894–2010). In the Palearctic, the oldest successfully DNA barcoded *Ph. issikii* specimen (obtained sequence length 408 bp) was a 162-year-old larva dissected from the leaf mine on *Tilia amurensis* from RFE (village Busse, Amur Oblast, the year 1859), sequence ID LMINH119-19. In the Nearctic, the oldest sequenced specimen (obtained sequence length 658 bp) was 127-year-old larva of *Ph. tiliacella* on *T. americana* from USA, Pennsylvania.

The length of recovered COI sequences ranged from 120 base pairs to 658 bp (Fig. 4). The sequence length for the archival insect specimens was negatively and significantly correlated with the specimen age (R^2^ = 0.32, N = 93, *P* < 0.05). Relatively long sequences (> 60% of the total length, i.e. the sequenced length > 400 bp) were obtained for 71 archival specimens having the age of 7–162 years (Fig. 4, the points in dashed rectangle) (Supplementary Table S4). Nine of these 71 specimens were over one century old (106–162-year-old): eight originated from the Palearctic and one from the Nearctic (Fig. 4, the points in gray cloud).

**Figure 4.**
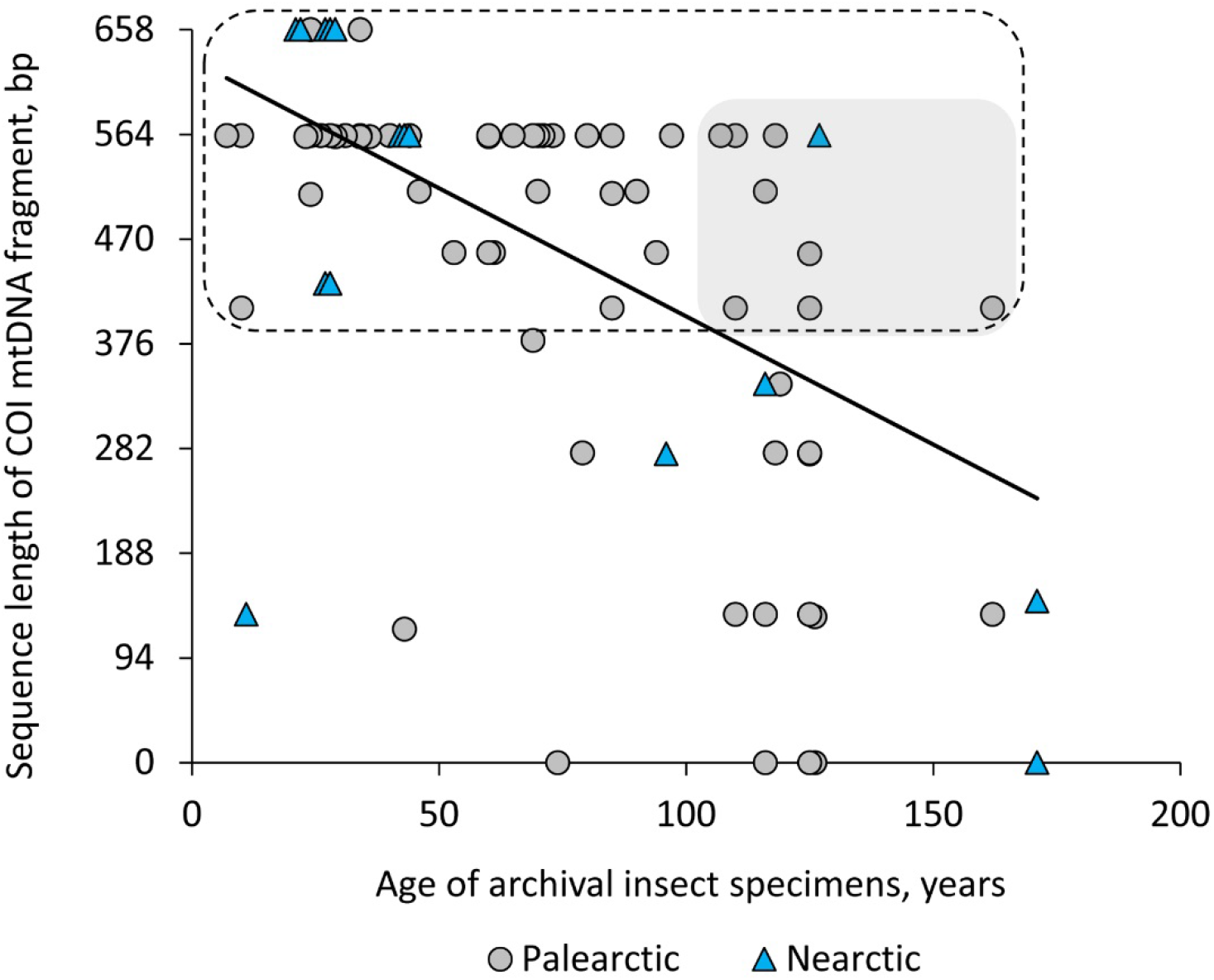
The relationship between the length of sequenced COI mtDNA fragment and the age of the archival *Phyllonorycter* specimens dissected from the mines in herbaria sampled in the Palearctic and Nearctic in 1850–2016. A linear regression is shown in the figure; y = –2.4x + 630, R^2^ = 0.32, N = 93, *P* < 0.05.

The 71 sequences represented five distinct clusters, each corresponding to a unique BIN (Fig. 5).

**Figure 5.**
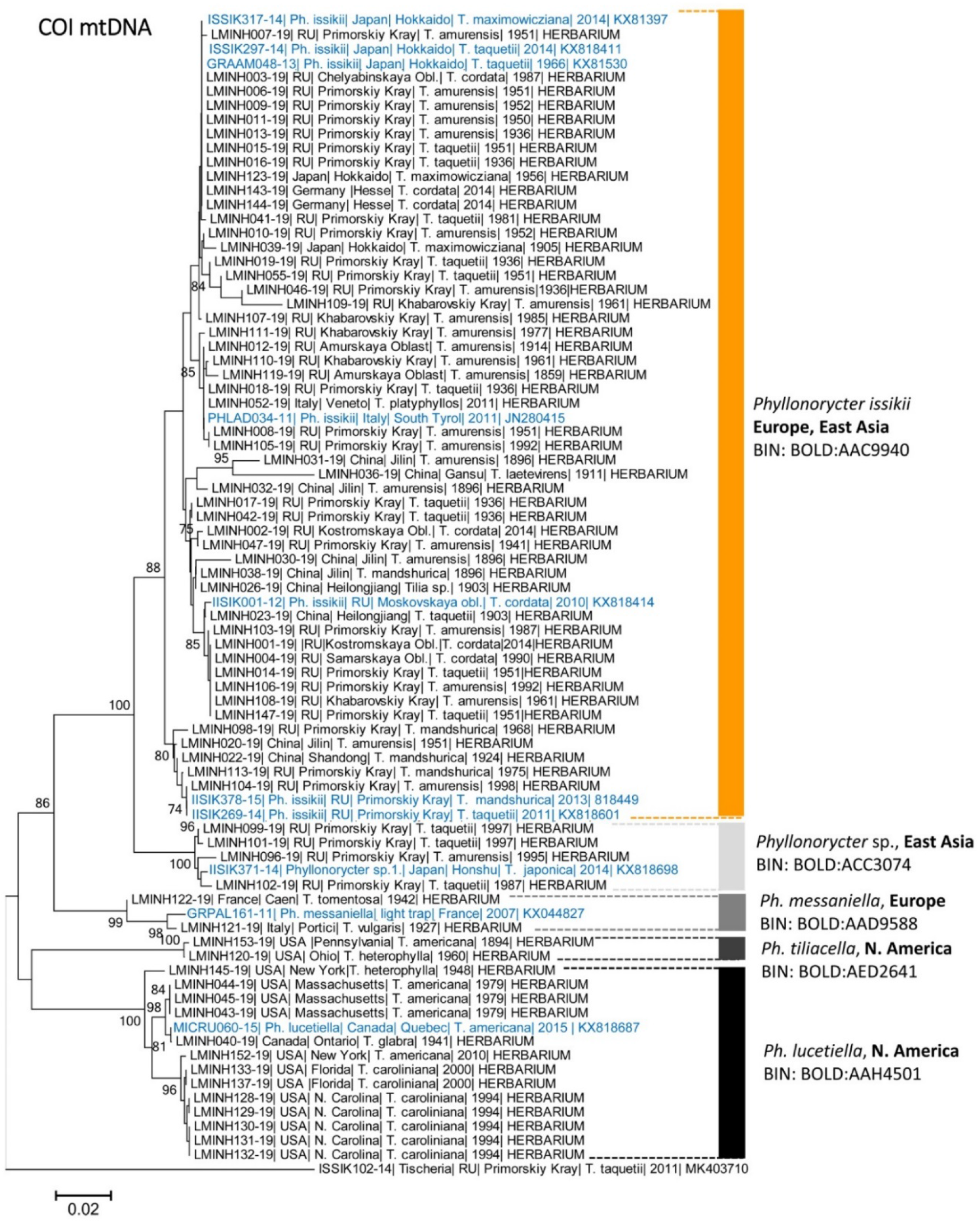
A maximum likelihood tree showing the proximity of the archival *Phyllonorycter* spp. archival specimens dissected from herbaria collected in the Palearctic and the Nearctic in 1859–2014. The tree was generated with the K2P nucleotide substitution model and bootstrap method (2500 iterations), *p* < 0.05. Each specimen is identified by its Process ID code | country | region | host plant | sampling year | genbank number (for modern sequences) or an indication of the source of the material (HERBARIUM) for archival specimens. Specimens in blue highlight reference sequences from modern range published in our previous study^20^. Each genetic cluster is specified by its Barcode Index Number (BIN) (given next to each cluster). Branch lengths are proportional to the number of substitutions per site.

Among them, two BINs with a Palearctic distribution were identified as *Ph. issikii* (number of specimens, N = 50, from 1859–1981) and *Ph. messaniella* (N = 2, from 1927 and 1942), and two BINs from the Nearctic were determined as *Ph. lucetiella* (N = 13, 1941–2010) and *Ph. tiliacella* (N = 2, 1894 and 1960). The fifth BIN was formed by an unidentified species of *Phyllonorycter* from RFE (N = 4, dated 1987–1997) (Fig. 5), a putative new species from the RFE and Japan^20^, showing a minimum pairwise distance of 4.79% with *P. issikii* (Table 2).

**Table 2.** Intra- and interspecific genetic divergences in DNA barcode fragments (COI mtDNA) in the archival specimens of lime-feeding *Phyllonorycter* spp. dissected from herbaria in the Palearctic and Nearctic^*^.

| Species, (realm)<sup>1</sup> | Species |  |  |  |  |
| --- | --- | --- | --- | --- | --- |
|  | *Ph. issikii* | *Ph. messaniella* | *Ph. tiliacella* | *Ph. lucetiella* | *Phyllonorycter* sp. |
| *Ph. issikii* (Palearctic) | [3.1] |  |  |  |  |
| *Ph. messaniella* (Palearctic) | 5.75 | [0.72] |  |  |  |
| *Ph. tiliacella* (Nearctic) | 9.69 | 10.48 | [0.18] |  |  |
| *Ph. lucetiella* (Nearctic) | 11.04 | 7.58 | 10.48 | [1.86] |  |
| *Phyllonorycter* sp. (Palearctic) | 4.79 | 6.11 | 13.14 | 11.55 | [1.06] |

\*Kimura two-parameter (K2P) distances (%); each species pair is provided with minimal pairwise distances; values in square brackets show maximal intraspecific distances. Number of analyzed archival specimens (N): *Ph. issikii* (50), *Ph. lucetiella* (13), *Ph. messaniella* (2), *Ph. tiliacella* (2), *Phyllonorycter* sp. (4).

The BIN of *Ph. issikii* contained 50 archival specimens: 42 from the East and 8 from the west of the Palearctic. In the East, the archival specimens were relatively old (40–162 years old) and originated from the RFE (32 specimens from 1859–1981), China (8 specimens from 1896–1924) and Japan (2 specimens from 1905–1956) (Fig. 5). On the other hand, in the West, the archival specimens of *Ph. issikii* were relatively recent (7–34 years old) and originated from Europe (2011–2015) and European part of Russia (1987–2015) (Fig. 5).

### Haplotype diversity and phylogeography of *Ph. Issikii*

Haplotype diversity in *Ph. issikii* past populations was significantly higher in the East (0.93) than in the West (0.65) (Mann-Whitney U test: U = 6, Z = – 0.75, *P* < 0.005).

Overall, 25 haplotypes were found among 50 sequenced archival specimens of *Ph. issikii* (Figs 6, 7, Table 3). All of them were detected in East Asia, including sex which were shared with the West (H1, H2, H8, H13, H22, and H23) (Fig. 6, Supplementary Table S5). The most common haplotypes were H1 (red) and H23 (green) (Figs 6, 7). Altogether they were found in 44% of all sequenced archival specimens, i.e. in 15 specimens (H1) and in 7 specimens (H23) out of 50 studied specimens. Notably, the haplotypes H1 and H23 dominate in the modern populations *Ph. issikii* and widely found both in the Eastern and Western Palearctic^20^. Based on the analysis of archival herbaria, the haplotypes H13 and H23 were for the first time detected in RFE and China, and the presence of the most distributed haplotype H1 was confirmed by historical material from Japan and RFE (Table 3) suggesting the contribution of Russia Far Eastern, Chinese and Japanese populations to the invasion of the moth westwards. Three haplotypes (H26, H28, H30) recorded in the archival *Ph. issikii* specimens from China and RFE have been already known from East Asia through sequencing the insect specimens sampled in nature in 21^st^ century, as per our recent study^20^ (Supplementary Table S5).

**Table 3.**
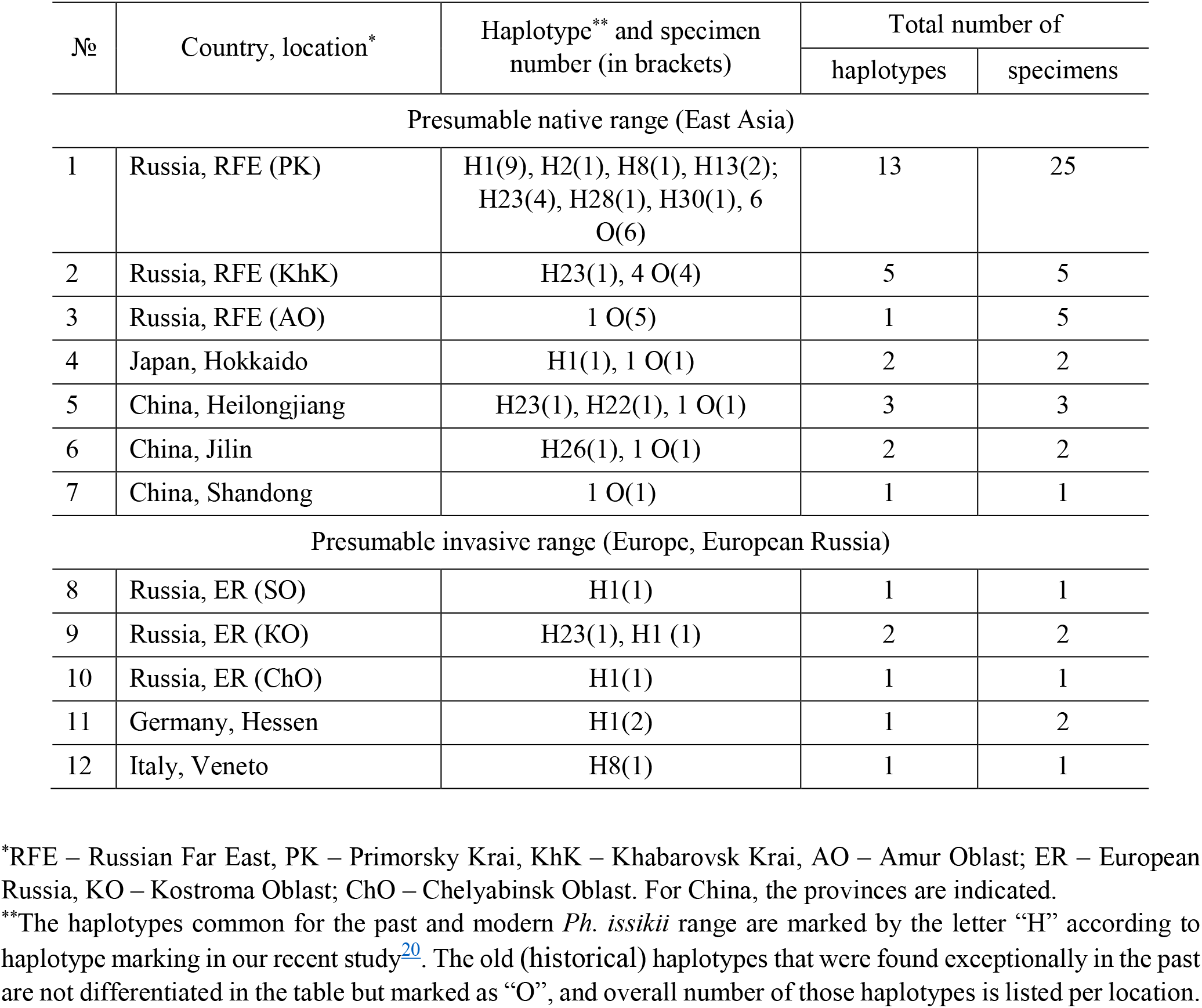
The haplotypes of *Phyllonorycter issikii* detected in the archival larvae and pupae that were dissected from the leaf mines on lime historical herbaria collected in the Palearctic; three haplotypes (H1, H8, H28) were shared between the East and the West.

**Figure 6.**
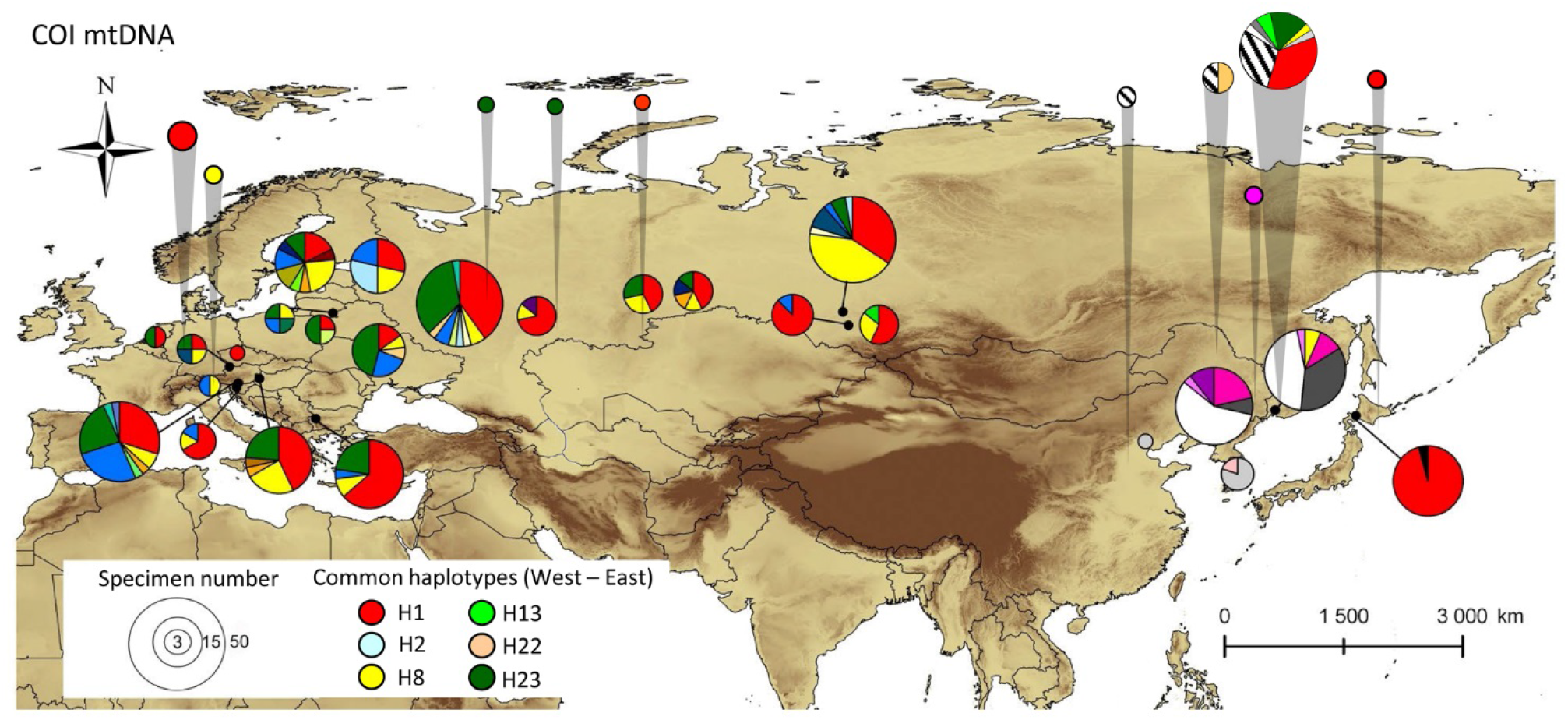
The distribution of historical and modern haplotypes of *Phyllonorycter issikii* in the Palearctic. The data on the haplotype diversity of *Ph. issikii* in the modern range modified from our previous study^20^. Each pie chart represents a country, except Russia where nearest locations are merged into one pie chat. The haplotypes from the historical area are indicated by gray callouts. Colors of pie charts refer to haplotypes found also in the modern specimens, as per^20^. The old (historical) haplotypes are counted altogether into one sector of a circle diagram with line shading (). The distribution of historical haplotypes by country can be found in Supplementary Tables S5, S6.

**Figure 7.**
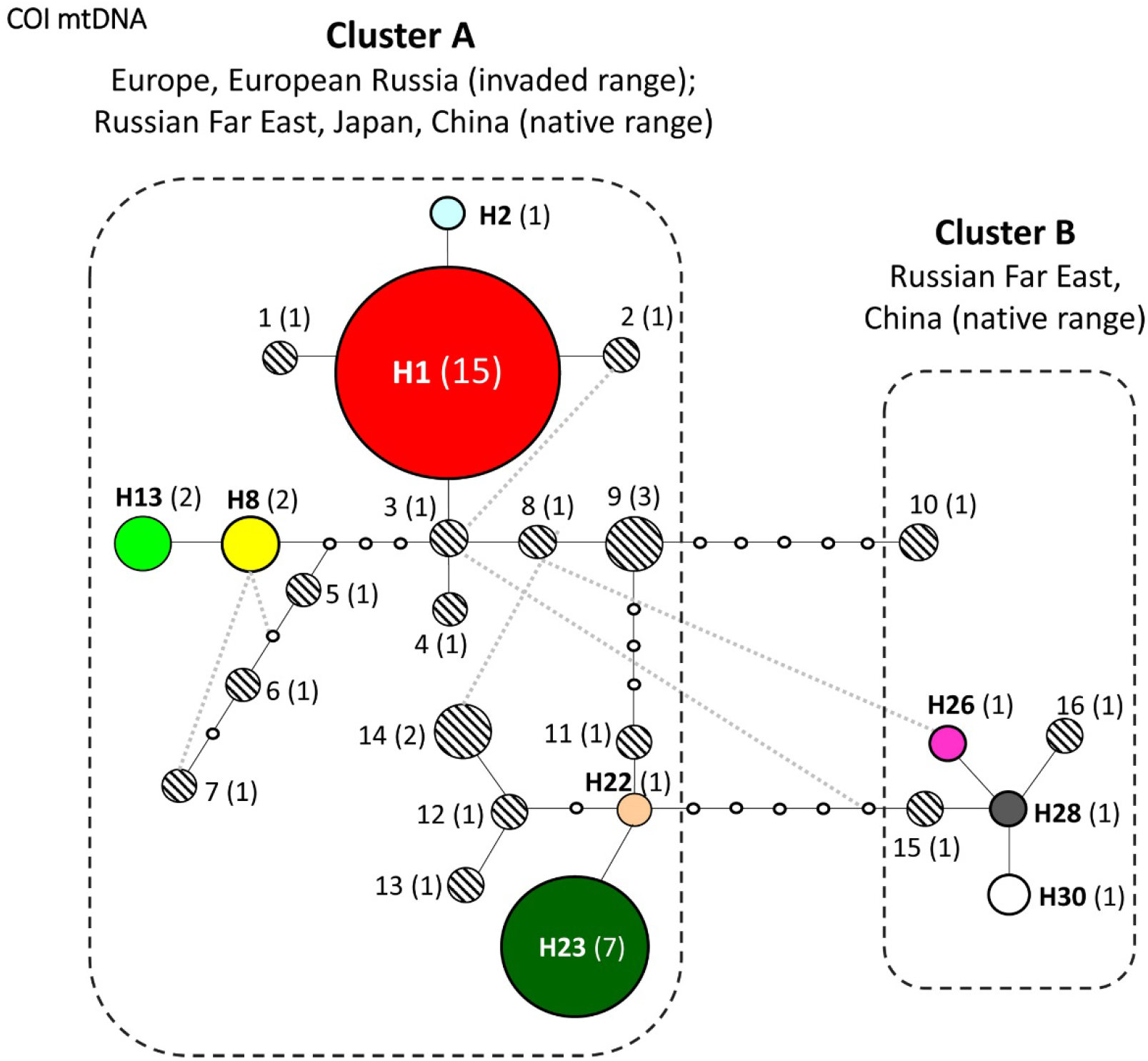
Median haplotype network of *Phyllonorycter issikii* in the Palearctic based on sequencing of archival specimens dissected from historical herbarium (1859–2014), *p* < 0.05. Different colors correspond to the haplotypes known in the modern *Ph. isskii* range^20^. These shared haplotypes are identically named, as follows: H1, H2, H8, H13, H22, H23, H26, H28, H30. Old (historical) haplotypes (documented for the first time from archival specimens) are indicated by numbers 1–16 next to the corresponding circles with shading. Empty tiny circles indicate intermediate missing haplotypes. The number of sequenced individuals is given in brackets next to each haplotype. Each line connecting the circles indicates one mutation step. Dotted rectangles indicate the two main geographically differentiated clusters. The geographical distribution of haplotypes is given in Supplementary Tables S5, S6.

In total, 16 out of 25 haplotypes detected in archival specimens from East Asia were old, i.e. they have not been found in the modern *Ph. issikii* populations (Fig. 7, Supplementary Table S6).

The reconstruction of the haplotype network supported the presence of two genetically differentiated clusters (A and B) in the past range of *Ph. issikii*, separated by 5 mutations (Fig. 7). Cluster A was formed by haplotypes found in both the western and eastern parts of the Palearctic, whereas Cluster B included exclusively haplotypes found in East Asia (Fig. 7).

## Discussion

The data from historical herbaria confirmed the hypothesis of *Ph. issikii* origin from East Asia and revealed the species distribution in the Russian Far East and China far prior to its description from Japan^14^. Furthermore, outbreaking levels in *Ph. issikii* populations in its native range were detected well before the species invasion into Western Palearctic. Indeed, herbarium specimens (*T. amurensis, T. taqueti*), from the RFE collected between 1934 and 1958 had up to 100% of leaves with mines (up to 13 mines per leaf). Prior to our study, no data was available in the literature recording significant damage by *Ph. issikii* in the native range (East Asia), except observation of an outbreaking density of *Ph. issikii* in Sapporo, Hokkaido (Japan) in 2002^20^. In contrast, outbreaking densities have been regularly recorded in Europe since the first discovery of the species in European Russia in 1985^19,24,25^.

Our study confirms that DNA preservation in archival larvae and pupae sealed in tissue of pressed leaves is sufficient for identification of century-old insect remains. Indeed, we were able to sequence the archival larvae and pupae dissected from 40-to-162-year-old herbaria obtaining sequence length of 120– 658 bp for 95% of all sequenced specimens (including those of 400–658 bp for 76% of specimens) and identify *Ph. issikii* in the sampling set. As expected, the obtained sequence length was negatively correlated with the age of insect specimens retrieved from mines in pressed lime specimens. A similar negative relationship was found for museum specimens of geometrid moths whose ages reached up to 157 years and which were DNA barcoded with Sanger sequencing^26^. The length of recovered sequences also showed the tendency to decrease with the age in DNA barcoded century-old Lepidopteran specimens^27,28^, with more encouraging results obtained with high-throughput sequencing compared to Sanger sequencing^27^.

Recent improvements in recovery of DNA barcodes from old and degraded samples enabled us also to distinguish the archival larvae of *Ph. issikii* and species alike, in particular to assign the mines in herbarized lime specimens (*T. tomentosa* and *T. cordata* f. *vulgaris*) sampled in England (1915–1987), France (1927), and Italy (1942) to another leaf mining moth, *Phyllonorycter messaniella*. It is a polyphagous species with the natural range in Europe and European Russia^29,30^. Developing on plants from several families, with oak as a main host (Fagaceae), in rare cases its mines can be found in Europe on limes, *Tilia* spp. (Malvaceae)^30,31^. Had we used morphology of the mines alone, we could have misdiagnosed this species as *Ph. issikii* and that would have significantly affected the dating of *Ph. issikii* invasion to Western Palearctic.

Using DNA-based identification tool, we showed lack of evidence of *Ph. issikii* introduction to the Nearctic in the past. Instead, we confirmed the presence of mines of two North American species in herbaria from Canada and the USA, namely *Ph. tiliacella* and *Ph. lucetiella*, previously known in these countries by literature^32^. Our study allowed to document the occurrence of these two species in several US states (see Supplementary Fig. S3) where they were not known^30^.

As expected, the analysis of archival *Ph. issikii* specimens revealed hidden haplotype diversity in the East, supporting the hypothesis of East Asia as the area of the moth’s origin. We discovered the haplotypes in the past *Ph. issikii* populations in Asia that had only been known in the species modern range from the invaded regions^20^. Furthermore, according to the archival herbaria, not only RFE and Japan, but also China were involved in the pest invasion westwards. To our big surprise, herbarium material and DNA sequencing analysis revealed broad presence of *Ph. issikii* in China suggesting that the occurrence of the species had been overlooked in China for decades. Interestingly, in China the species was not known before our first record in Tianjin^20^. Herbarium study revealed archival mines in 15 (!) different Chinese provinces, mostly in northeast and central parts of the country. The archival mines from Southern China provinces (refer to Indomalaya) did not differ from those of *Ph. issikii*. However, the species identity could not be proven as no larvae or pupae were found in the mines in the herbarium specimens originating from these provinces for morphological or DNA-based identification. Thus, further work is needed to confirm the presence of *Ph. issikii* in Southern China.

*Ph. issikii* is currently known widely in Europe and European Russia, almost everywhere where limes are present^18,19,20^. According to the herbarium and DNA barcoding data, *Ph. issikii* occurred in Western Palearctic in the 1980s. The earliest archival mine found in Europe dates back to 1987 in European Russia (Fig. 8). These results agree with *Ph. issikii* documentation in Western Palearctic in literature: the first finding of the species outside East Asia referred to Moscow and dated back to 1985^33^ (Fig. 8). Two years after, a foci of *Ph. issikii* was recorded in Voronezh, 500 km south-west of Moscow^25^ (see also Fig. 8), whereas in the next two decades it was recorded almost everywhere in European Russia^19^.

**Figure 8.**
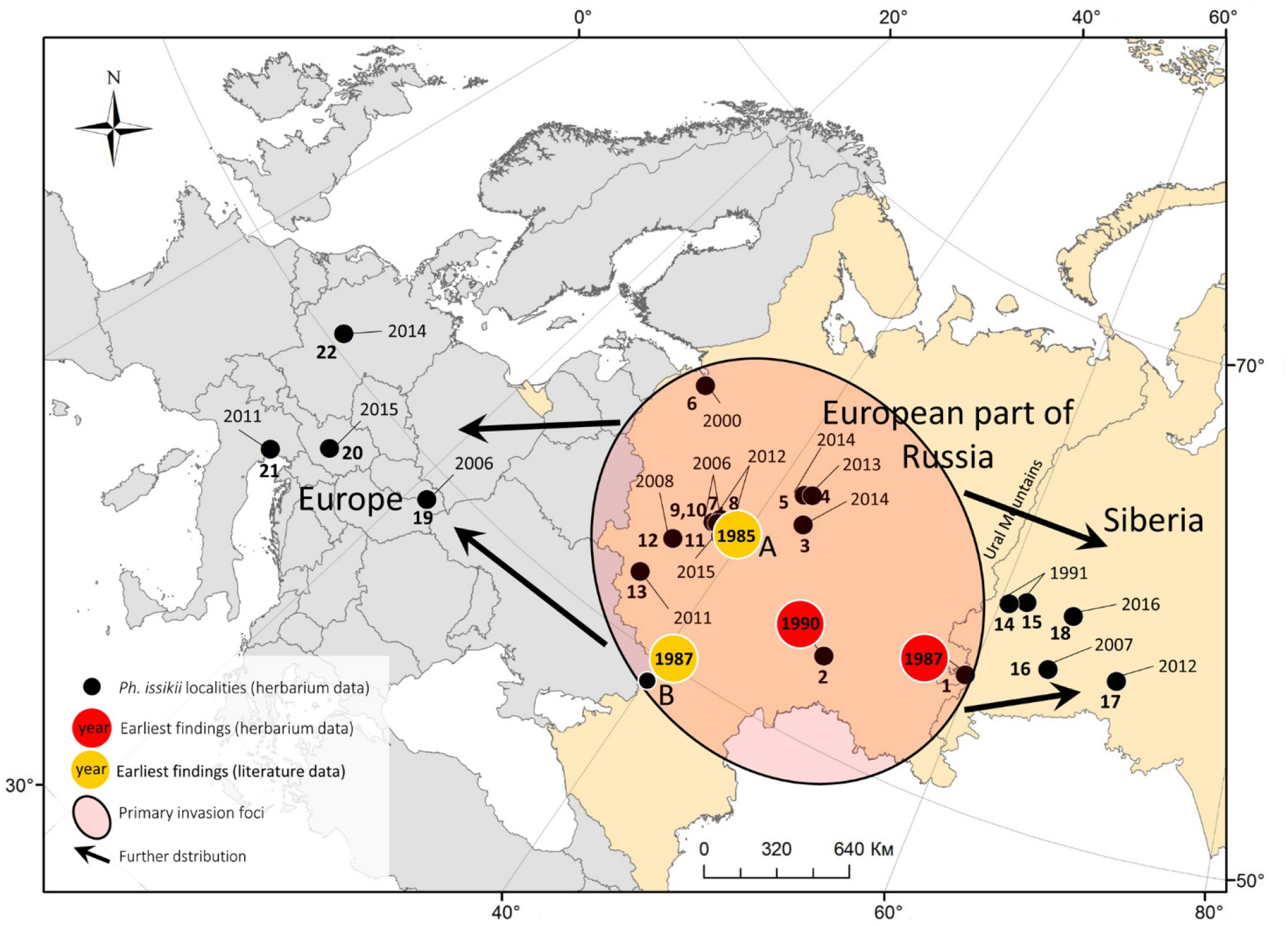
The suggested scenario of *Phyllonorycter issikii* invasion in Western Palearctic based on archival data from the historical herbaria and published records. The dots numbers (1–22) indicate the regions or countries where mines were found in herbaria. Next to the dots, the year of collection of herbarium specimens with *Ph. issikii* mines is indicated: 1–18 – Russia (1 – Chelyabinsk, 2 – Samara, 3–5 – Kostroma, 6 – Leningrad, 7–11 – Moscow, 12 – Kaluga, 13 – Kursk, 14–15 – Sverdlovsk, 16–18 – Tyumen Oblasts), 19 – Slovakia, 20 – Austria (Styria), 21 – Italy (Veneto), 22 – Germany (Hessen). Black arrows show possible directions of *Ph. issikii* expansion from the primary foci.

The dates of the first *Ph. issikii* records in the literature (1985 in Moscow) and those obtained in the present herbarium study (1987 in Zlatoust) are very similar (Fig. 8) but the distance between these two locations, situated in the opposite parts of European Russia, i.e. in its western and eastern parts respectively, is considerable (about 1400 km). The findings of *Ph. issikii* in a near time in so distant locations may suggest that, by the time of the species revelation, it had already spread across significant territory of European Russia that can also be a result of *Ph. issikii* multiple introductions from Eastern Palearctic^20^.

An unintentional introduction of *Ph. issikii* into European Russia could have happened after the Second World War when green landscaping in damaged cities and settlements required fast and effective restoration and greening programs^34,35^. At that period, botanical gardens were intensively created in European Russia, with numerous introductions of woody plant species from the East^36^. Thus, *Ph. issikii* may have escaped East Asia with its host plants, in particular *T. amurensis* and *T. mandshurica*, which have a natural range in East Asia and are nowadays known in culture across most of European Russia^37^. However, we were unable to test this hypothesis due to what herbarium specimens were collected scarcely across Europe, European Russia, and Siberia in 1940–1970s.

Although we were unable to find earlier evidences of *Ph. issikii* invasion westward despite extensive surveys of herbaria, our study clearly shows that in the West, the primary invasion foci occurred in European Russia, from where the moth started expanding westward to other European countries and eastward to Western Siberia (Fig. 8). These findings concur with the documentation of the moth’s distribution in these directions, as documented in the literature^19^.

To conclude, the survey of the large herbaria collected during the last 253 years in the northern hemisphere allowed us to clarify the invasion history of the lime leaf mining micromoth, *Phyllonorycter issikii* and confirm its East Asian origin and the invasive status in the west of the Palearctic. The findings of the characteristic leaf mines in herbaria and molecular data recovered from larvae and pupae found in the mines in pressed leaves significantly improved our knowledge about past range of the leaf miner in East Asia, and identified its trophic associations with East Asian limes. Moreover, it pointed at the contribution of *Ph. issikii* populations from the Russian Far East, Japan and China to the invasion westward of the Palearctic and confirmed no occurrence of the species in the Nearctic. Finally, our study highlighted the importance of historical herbaria in studying past distribution and diversity of endophagous folivore organisms, including pestiferous and invasive species. As such, sampling and storing herbarium collection, including plant vouchers carrying any type of damage, and their wider use can be especially beneficial for invasive ecology research.

## Methods

### Studied herbarium collections

To reconstruct the past range and clarify the invasion history of *Ph. issikii*, we examined historical herbaria of *Tilia* spp. collected in the last 253 years (1764–2016) in the Palearctic, deposited in 20 museums and botanical gardens across Eurasia (Supplementary Table S7). Additionally the herbarium collected in the last 109 years (1818–2008) in the Nearctic, where limes are also distributed^38^, was examined to check for *Ph. issikii* introduction in the past. So far, the moth has not been documented from the Nearctic^39^.

In total, we examined 15,686 herbarium specimens of *Tilia* of which 15,003 specimens (96%) originated from the Palearctic and 683 (4%) from the Nearctic (Supplementary Fig. S4), altogether accounting for about 1.4 million lime leaves. The majority of herbarium specimens from the Palearctic (11,454, i.e. 76% of all studied specimens from this region) came from the putative invasive range of *Ph. issikii* (i.e. from Europe, European Russia and Western Siberia). In Europe alone, the specimens originated from 30 different countries, altogether accounting for 8444 specimens (56%) (Supplementary Fig. S4). Nearly in all these countries, *Ph. issikii* was known by literature records^18,19,20,40^, except in the Scandinavian countries and England, where *P. issikii* has not been found yet^30^. In European Russia and Western Siberia, the herbarium was represented by 2696 (18%) and 314 lime specimens (2%) respectively (Supplementary Fig. S4). The other 3,549 herbarium specimens (24% of all studied material in the Palearctic) originated from the putative native range, East Asia, in particular from RFE (2,202 specimens, 15%), China (940 specimens, 6.2%), Japan (339 specimens, 2.3%), and Korea (68 specimens, 0.5%) (Supplementary Fig. S4). *Ph. issikii* was historically known from these regions/countries (with the type locality in Japan)^14,15,16^, except from China, where it was recorded for the first time in 2015^20^.

In the Nearctic, 638 out of 683 herbarium specimens originated from North America from 16 US states (604 specimens, i.e. 88% of all specimens studied in the Nearctic) and from two Canadian provinces (34 specimens, i.e. 5%) (Supplementary Fig. S4). The other 45 herbarium specimens (7%) originated from Northern Mexico (Supplementary Fig. S4).

### Surveyed lime species

Given the ability of *Ph. issikii* to develop on different limes^14,20,40^, specimens of all representatives of the genus *Tilia* stored in herbaria were examined. Altogether they accounted for 117 lime species and more than 80 hybrids across the Palearctic and Nearctic (Supplementary Table S8). Bearing in mind that the taxonomy of limes has been revised several times^41,42^, some species indicated on the labels of herbarium specimens have been synonymized by botanists since the collection date. However, in order to keep the historical aspect and avoid misinterpretation of the primary identifications, we refer to the lime species names as they were originally stated on the vouchers.

### Examination of herbarium specimens

The herbarium specimens were carefully examined for the presence of the typical *Phyllonorycter* mines, i.e. whitish oval flat or tentiform blotches on leaf lamina (Fig. 9). Only the mines of a diameter 3–5 mm onward were taken into consideration (by the end of development, the diameter of *Ph. issikii* mines may reach 15 mm). The mines of a smaller size are, in general, difficult to spot on pressed leaves and to identify reliably.

**Figure 9.**
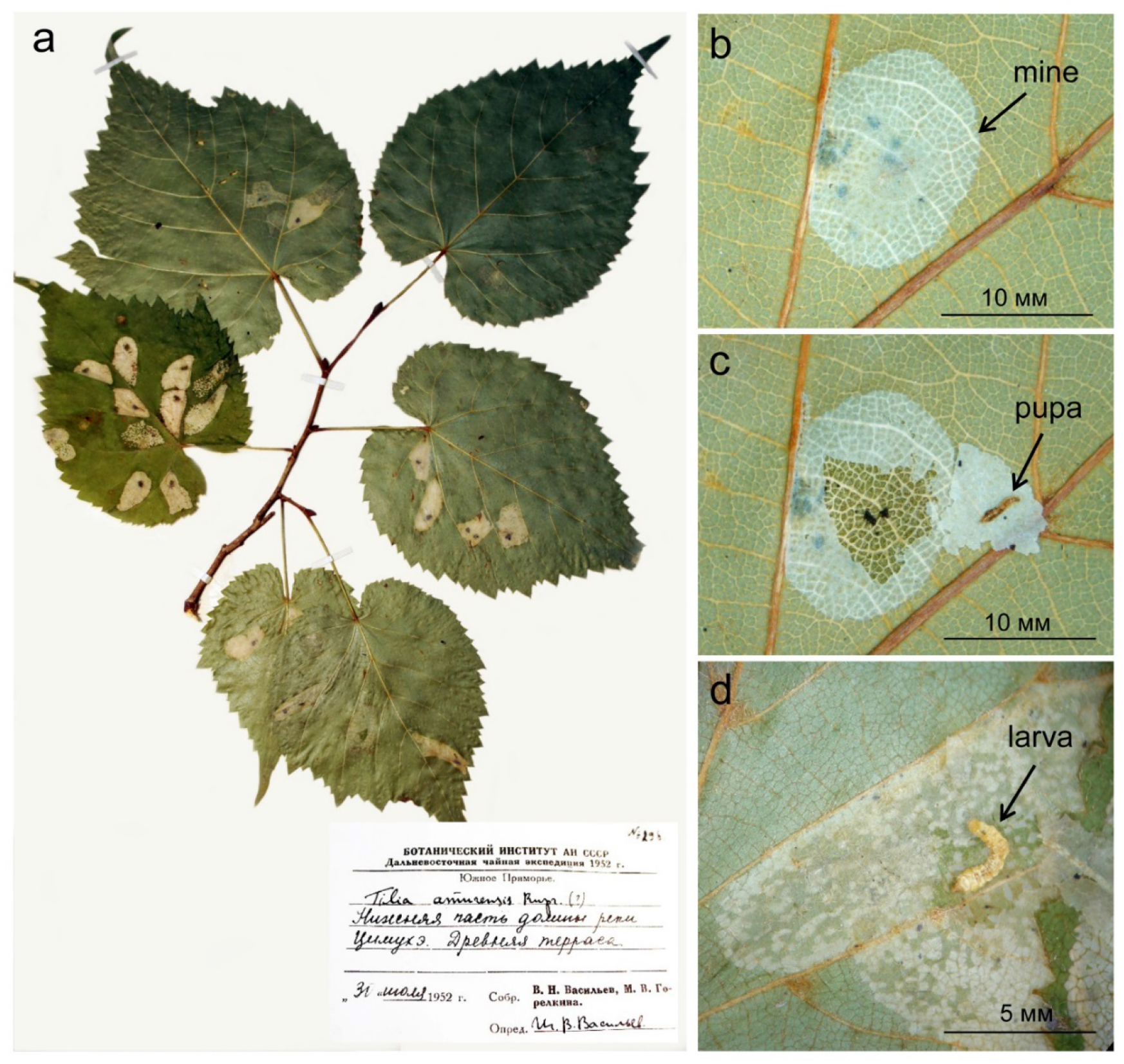
An example of herbarium specimen with *Phyllonorycter issikii* mines (**a**), the mine (**b**), opened mine with the pupa (**c**), opened mine with the larva (**d**). The data from the label at the herbarium specimen originating from the Russian Far East: № 296, Botanical institute of the Academy of Sciences of USSR, Far Eastern Tea Expedition of the year 1952; South Primorye, *Tilia amurensis* Rupr., the lower part the valley of the river Tsimukhe, ancient terrace; 31.VII.1952 coll; V. N. Vasilyev, M. V. Gorelkina coll.; I. V. Vasilyev det.

The pressed leaves were examined from the lower side where *Ph. issikii* lay eggs and make blotch mines^14,18^. The total number of leaves in a herbarium specimen, the number of leaves with the mines, and the number of mines on the leaves were counted.

Randomly selected mines were opened to sample late instar larvae or pupae for further analyses (Fig. 9). The mines were opened by making an incision in the epidermis in the area where larvae or pupae were spotted (Fig. 9). To avoid contamination, the forceps used for picking insects were properly cleaned with 95% ethanol solution after each manipulation. The sampled larvae and pupae (93 individuals) were placed individually into 2ml microtubes with hermetically sealed lids (Axygen, USA), labeled and stored in a freezer at –20°C prior for morphologic (pupae) and molecular genetic (larvae and pupae) analyses. Additionally, 17 pupal exuvia were sampled for morphological identification.

### Morphological diagnostics

Pupae and pupal exuvia were examined at magnification × 20–40 using ZEISS Stemi DV4 binocular (Germany); pupal cremaster was studied under the magnification × 60 using the microscope Olympus CX21 (Japan). Based on the mine characters (i.e. the position of mines on the leaf surface, presence/absence of folds on epidermis covering mine, frass pattern) and the sculpture of pupal cremaster^31,32,43^, the mines were attributed to lime-feeding *Phyllonorycter* species known in the Palearctic and Nearctic (Supplementary Table S9). The typical oval mines, mostly situated between the secondary veins on the lower epidermis, with no folds on epidermis covering mine^31^, inside which neither larvae nor pupae were found from the localities where *Ph. issikii* is known in the Palearctic were attributed to the latter.

### Molecular genetic analysis

Archival larvae and pupae dissected from the mines on pressed leaves (overall 93 specimens) were analyzed using DNA barcoding paired with third generation sequencing to confirm species identity, determine historic haplotypes, overview past range, find early evidences of *Ph. issikii* invasion in the Palearctic, and check for the species presence in the Nearctic back in time. Nondestructive approach was used to save the vouchers. In the archive insects, up to 11 overlapping fragments (ranging from 150 to 230 nucleotides) of the mtDNA COI gene region (658 bp) were sequenced. DNA extraction, amplification, and sequencing were performed at the Canadian Center for DNA-barcoding at the University of Guelph (Canada). The molecular protocols from a previous study^23^ were adapted for single molecular real-time sequencing (SMRT) on Sequel platform (Pacific Biosciences)^27^. For the assembly of nucleotide contigs, multiple fragments were aligned to the reference sequence using the Codon Code Aligner V.3.7.1 software (CodonCode Corporation).

### Data analysis

The Spearman’s rank correlation (R) was used to assess sequencing success, i.e. the relationship between the length of the sequenced fragment (maximal of 658 bp) and the age of the archival specimens, (*p* <0.05). In further analyses only sequences with the length of at least 300 bp were used; shorter sequences with missing diagnostic sites were excluded. Archival specimens were identified by their DNA barcodes to the species level using BOLD identification engine^44^. The phylogenetic tree analyzing the relatedness of the archival specimens was built using the Maximum likelihood estimation algorithm, the Kimura’s two-parameter (K2P) model and the bootstrap with 2500 iterations. In the phylogenetic tree, the topology of the basal branches was considered robust at a value ≥ 70. Phylogenetic analysis was run in Mega X^45^. The haplotype diversity was computed as follows, H = (N / (n – 1)) × (1 – ∑x_i_^2^), where x_i_ is the relative haplotype frequency of each haplotype in the sample and N is the sample size^46^.

To determine the haplotypes, the sequences of the archival *Ph. issikii* specimens were analyzed in the total alignment with the pool of the DNA barcodes previously accumulated in the study of 377 *Ph. issikii* individuals in the modern species range in the Palearctic, dx.doi.org/10.5883/DS-TILIAPHY^20^. For sequences with different lengths, only the length aligned with compared sequences was taken into the consideration for distance calculation and haplotype determination. The haplotypes identified in archival specimens, that were identical to those occurring in the modern *Ph. issikii* range, as determined in our previous study^20^, were denoted the same haplotype name (for example, H2 – “Haplotype 2”, common for modern and historical ranges) and shaded by the same color on the haplotype network. The haplotypes identified in the archival specimens that differed from those known in the *Ph. issikii* modern range were considered as “old” haplotypes, i.e. present in the past but absent in the moth’s modern range. The median-joining haplotype network was constructed in TCS 1.21 implementing a statistical parsimony algorithm^47^.

The geographic distribution of COI haplotypes was mapped using ArcGIS 9.3^48^. The historical range of *Phyllonorycter* species was illustrated in accordance with the finding time scale (in a 10-year interval). The *Ph. issikii* distribution scenario in Western Palearctic was generated based on the dating records of the first occurrence of *Ph. issikii* in different locations as per published data^18,19,49^ and the herbarium data analyses performed in the present study.

In the Palearctic, the number of herbarium specimens carrying *Phyllonorycter* mines was analyzed for the putative native range (East Asia), that we referred to the Eastern Palearctic, versus the putative invaded range (European countries, European Russian and Western Siberia), that we referred to Western Palearctic. The average number of leaves with the mines per herbarium specimen was calculated and compared between these two macroregions. We also compare the East and the West by the dating of the mines in historical herbaria (i.e. distribution of records across time scale). The Mann– Whitney nonparametric U-test, allowing analyzing small and unequal sampling sets, was used for data comparison (STATISTICA 12.6 software, Stat Soft. Inc., USA).

## Ethics statement

The herbaria surveys were carried out in respect to the Institutional and International ethical guidelines. The protocol of archival larvae and pupae sampling from herbarized leaves was approved by the herbarium curators in all visited herbarium depositaria.

## Data availability

The specimen data and the genetic dataset (together with sequence records, and GenBank accession numbers) generated during the study are publicly available from BOLD (The Barcode of Life Data System) using the link: (doi, pending).

## Acknowledgments

We greatly thank the curators and staff at herbarium depositaria in Eurasia: Vanessa Invernón (the herbarium code P at National Museum of Natural History, Paris), Sally Dawson (R at Royal Botanic Gardens, Kew), Jovita Yesilyurt (BM at Natural History Museum, London), Lesley Scott (E at Royal Botanic Garden Edinburgh, Edinburgh), Ashleigh Whiffin (National Museums Collection Center, Edinburgh), Roxali Bijmoer (L at Naturalis Biodiversity Center, Leiden), Chiara Nepi (FI at Natural History Museum, Florence), Agnese Tilia (RO at Sapienza University of Rome, Rome), Nicolas Fumeaux (G at Conservatory and Botanical Garden of the city of Geneva, Geneva), Alessia Guggisberg (ZT at Botanical Garden Zürich, Zurich), Peter Hein and Sarah Bollendorff (B at Berlin Botanic Garden and Botanical Museum, Berlin), Christian Bräuchler (W at Natural History Museum, Vienna), Larisa Orlova (LECB at Komarov Botanical Institute of the Russian Academy of Sciences, Saint Petersburg), Maria Nosova, Mikhail Ignatov and Nina Stepanova (MHA at Principle Botanical Garden RAS, Moscow), Alexey Seregin (MW at Moscow State University, Moscow), Irina Gureeva (TK at Tomsk State University, Tomsk), Natalia Kovtonyuk (NS at Central Siberian Botanical Garden SB RAS, Novosibirsk), Valentina Verkholat (VBGI at Botanic Garden–Institute FEE RAS, Vladivostok), Zoya Kozhevnikova and Valentin Yakubov (VLA at Federal Scientific Center of the East Asia Terrestrial Biodiversity FEB RAS, Vladivostok), Irina Goncharova (KRF at Sukachev Institute of Forest SB RAS, Krasnoyarsk) for giving us an access to herbarium collections. Thanks to David Lees (London), Erik J. van Nieukerken (Leiden), Marina Fonti (Zurich), Lidia Seraya, Jeanna Agafonova (Moscow), Dmitry Musolin (Saint Petersburg), Svetlana Krivets (Tomsk), Maria Tomoshevich (Novosibirsk), Margarita Ponomarenko (Vladivostok), and David Lees (England) for helping with herbarium contacts and/or for their great hospitality during the work of N.I.K. in herbarium depositaria. Thanks to Jean-Francois Landry (Canada) and Charley Eiseman (USA) for consulting on morphology of mines and pupae of *Phyllonorycter lucetiella* and *Ph. tiliacella* and for sharing the photographs of mines and pupal cremaster with us. Thanks to Ulf Buentgen (Cambridge), Marc Kenis (Switzerland), Alain Roques (France), Yuri Baranchikov (Krasnoyarsk, Russia) for fruitful discussions and useful advices, Irina Mikhailova (Krasnoyarsk) for helping with mapping. We also thank Norman Monkhouse, Liuqiong Lu, and Sean Prosser at the Canadian Centre for DNA Barcoding (University of Guelph, Canada) for technical support.

This work was funded by Le Studium (France), Short Term Scientific Mission (STSM) within COST Action FP1401 “Global Warning” [herbarium survey in Europe], the Russian Foundation for Basic Research (No. 19-04-01029-a) [herbarium survey in Russia, molecular genetic data analysis], and the basic project of Sukachev Institute of Forest SB RAS (project No. 0287-2021-0011) [morphological analysis]. The DNA barcoding analysis was also supported in part by the “Food From Thought (FFT) Program” and Canada First Research Excellence Fund (CFREF).

This article is dedicated to the memory of the late Tosio Kumata (Japan) for his significant contribution to the knowledge of East Asian Gracillariidae, in particular, for discovering and describing *Phyllonorycter issikii*, as well as for fruitful discussions on this species during our (N.I.K., C.L.V.) visit to Japan.

## Author contributions

N.I.K. and C.L.V. designed the study, N.I.K. surveyed herbaria, sampled insect specimens, analyzed data, N.I.K., C.L.V., E.V.Z. ran molecular genetic analysis. The manuscript was written by N.I.K. and reviewed by C.L.V. and E.V.Z.

## Additional information

### Supplementary Information

The online version contains supplementary material available at https://doi.org/XXX (pending)

## References

1. Lang, P. L. M., Willems, F. M., Scheepens, J. F., Burbano, H. A., Bossdorf, O. Using herbaria to study global environmental change. New Phytologist 221, 110–122 (2018).

2. Heberling, J. M., Prather, L. A., Tonsor, S. J. The changing uses of herbarium data in an era of global change: an overview using automated content analysis. BioScience 69, 812–822 (2019).

3. Abbott, I., Wills, A., Burbidge, T. Historical incidence of Perthida leaf miner species (Lepidoptera) in southwest Western Australia based on herbarium specimens. Australian J. Ecol. 24, 144–150 (1999).

4. Lees, D. C. et al. Tracking origins of invasive herbivores using herbaria and archival DNA: the case of the horse-chestnut leaf miner. Front. Ecol. Environ. 9, 322–328 (2011).

5. Meineke, E. K., Dunn, R. R., Sexton, J. O., Frank, S. D. Urban warming drives insect pest abundance on street trees. PLoS ONE 8, e59687 (2013).

6. Farr, J. D. Herbarium specimens provide historical evidence of Cardiaspina jerramungae (Hemiptera: Psylloidea, Aphalaridae) outbreaks on Eucalyptus occidentalis in the Lower Great Southern of Western Australia. Aust. Entomol. 59, 167–177 (2020).

7. Staats, M. et al. Genomic treasure troves: complete genome sequencing of herbarium and insect museum specimens. PLoS ONE 8, e69189 (2013).

8. Antunes, P., Schamp, B. Constructing standard invasion curves from herbarium data – towards increased predictability of plant invasions invasive plant science and management. Invasive Plant Sci. Manag. 10, 293–303 (2017).

9. Beaulieu, C., Lavoie, C., Proulx, R. Bookkeeping of insect herbivory trends in herbarium specimens of purple loosestrife (Lythrum salicaria). Phil. Trans. R. Soc. B 374, 20170398 (2018).

10. Skvortsov, A. K. Herbarium. Methodology and technique manual (Moscow, Nauka, 1977). [in Russian]

11. Meineke, E. K., Tomasi, C., Yuan, S., Pryer, K. M. Applying machine learning to investigate long-term insect–plant interactions preserved on digitized herbarium specimens. Appl. Plant Sci. 8, e11369 (2020).

12. Kobayashi, S., Johns C. A., Kawahara, A. Y. Revision of the Hawaiian endemic leaf-mining moth genus Philodoria Walsingham (Lepidoptera: Gracillariidae): its conservation status, host plants and descriptions of thirteen new species. Zootaxa 9444, 1–175 (2021).

13. Hembry, D. Putative insect extinction on the deforested Island of Mangareva (Gambier Archipelago, French Polynesia). Pacific Science 67, 553–560 (2013).

14. Kumata, T. Taxonomic studies on the Lithocolletinae of Japan. Insecta Matsumurana 25, 53–90 (1963).

15. Ermolaev, V. P. A review of the fauna and ecology of miner-moths (Lepidoptera, Gracillariidae) of the Primorye Territory. The Fauna of Insects of the Far East 70, 98–116 (1977). [in Russian]

16. Kumata, T., Kuroko, H., Park, K. Some Korean species of the subfamily Lithocolletinae Gracillariidae, Lepidoptera. Korean J. Plant Protection 22, 213–227 (1983).

17. Kim, D.-S., Byun, B.-K. Taxonomic review of the genus Phyllonorycter Hübner (Lepidoptera: Gracillariidae) in Korea. J. Asia Pacific Entomol. 20, 1377–1386 (2017).

18. Šefrová, H. Phyllonorycter issikii (Kumata, 1963) – bionomics, ecological impact and spread in Europe (Lepidoptera, Gracillariidae). Acta Universitatis Agriculturae et Silviculturae Mendelianae Brunensis 50, 99–104 (2002).

19. Ermolaev, V. I., Rubleva, E. A. History, rate and factors of invasion of lime leafminer Phyllonorycter issikii (Kumata, 1963) (Lepidoptera, Gracillariidae) in Eurasia. Rus. J. Biol. Invasions, 1, 2 –19 (2017). [in Russian]

20. Kirichenko, N. et al. From east to west across the Palearctic: phylogeography of the invasive lime leaf miner Phyllonorycter issikii (Lepidoptera: Gracillariidae) and discovery of a putative new cryptic species in East Asia. PLoS ONE 12, e0171104 (2017).

21. Kirichenko, N., Augustin, S., Kenis, M. Invasive leafminers on woody plants: a global review of pathways, impact and management. J. Pest Sci. 92, 93–106 (2019).

22. Lopez-Vaamonde, C., et al. Evaluating DNA barcoding for species identification and discovery in European Gracillariid moths. Front. Ecol. Evol. 9, 626752 (2021).

23. Prosser, S., Dewaard, J., Miller, S., Hebert P. DNA barcodes from century-old type specimens using next generation sequencing. Mol. Ecol. Res. 16, 487–497 (2016).

24. Kozlov, M. V. Leaf-mining moth is a pest of Limes. Plant protection and quarantine 4, 46 (1991). [in Russian].

25. Šefrová, H. Invasions of Lithocolletinae species in Europe – causes, kinds, limits and ecological impact (Lepidoptera, Gracillariidae). Ekologia Bratislava 22, 132–142 (2003).

26. Strutzenberger, P., Brehm, G., Fiedler, K. DNA barcode sequencing from old type specimens as a tool in taxonomy: a case study in the diverse genus Eois (Lepidoptera: Geometridae). PLoS ONE. 7, e49710 (2012).

27. D’Ercole, J., Prosser, S., Hebert, P. A SMRT approach for targeted amplicon sequencing of museum specimens (Lepidoptera)-patterns of nucleotide misincorporation. PeerJ 9, e10420 (2021).

28. Hebert, P.D.N. et al. A DNA ‘barcode blitz’: rapid digitization and sequencing of a natural history collection. PLoS One 8, e68535 (2013).

29. Baryshnikova, S. V. Family Gracillariidae in Catalogue of Lepidoptera of Russia (ed. Sinev, S.Yu.) 36–43 (St. Petersburg: Zoological Institute RAS, 2nd ed., 2019). Baryshnikova, S.V., Dubatolov, V.V. (2007) To knowledge of small moths (Microlepidoptera) of the Nature Reserve “Bolshekhekhtsirskii” (Khabarovsk District). 2nd report. Bucculatricidae, Gracillariidae, Lyonetiidae. *Animal world of the Far East* **6**, 47–50 (2007). [in Russian]

30. De Prins J., De Prins W. Global taxonomic database of Gracillariidae (Lepidoptera). Belgian Biodiversity Platform BELSPO. http://www.gracillariidae.net/ (2021).

31. Ellis, W. Plant parasites of Europe: leafminers, gallers and fungi. The Netherlands. http://bladmineerders.nl/ (2021).

32. Eiseman, C. Leaf miners of North America [electronic source] 1st Edition (June version). World Press. http://charleyeiseman.com/leafminers/ (2019).

33. Bednova, O. V., Belov, D. A. The lime leaf miner (Lepidoptera, Gracillariidae) in green areas of Moscow and Moscow region. Bull. Moscow State For. Univ. 2, 172–177 (1999). [in Russian]

34. Lebedev, G. I. Green building in the cities of the USSR (Moscow-Leningrad: Publishing house of the Ministry of communal services of the USSR, 1948). [in Russian]

35. Protopopova, E. N. The recommendations for landscaping in cities and settlements in Central Siberia (Krasnoyarsk: Krasnoyarskiy Rabochiy, 1972). [in Russian]

36. Lapin, P. I., Kalutsky, K. K., Kalutskaya, O. N. Introduction of forest species (ed. Ogorodnikova L. M.) 224 p. (Moscow, Forest Industry, 1979). [in Russian]

37. Vasilyev, I. V., Sokolova, Ya. A. Limes – Tiliaceae Juss in Trees and bushes of the USSR (Moscow-Leningrad: Publishing house of the Academy of Sciences of the USSR) 659–727 (1958). [in Russian]

38. Pigott, C. D. Lime-trees and Basswoods. A biological monograph of the genus Tilia (New York: Cambridge University Press, 2012).

39. Leaf blotch miner moth Phyllonorycter issikii (Kumata). Center for invasive species and ecosystem health. University of Georgia, USDA, USA. https://www.invasive.org/–browse/subinfo.cfm?sub=12059 (2021).

40. Ermolaev, I. V., Rubleva, E. A., Rysin, S. L., Ermolaeva, M. V. Forage plants of lime leaf miner Phyllonorycter issikii (Kumata, 1963) (Lepidoptera, Gracillariidae). Rus. J. Biol. Invasions 2, 2–13 (2018). [in Russian]

41. Cronquist, A. The evolution and classification of flowering plants (Boston: Houghton Mufflin Company, 1968).

42. Koropachinsky, I. Yu., Vstovskaya, T. N. Wood plants of the Asian part of Russia (Academic Publishing House GEO, Novosibirsk, 2nd Edition, 2012). [in Russian]

43. Gregor, F., Patočka, J. Die Puppen der mitteleuropaischen Lithocolletinae. Mitteilungen des internationalen entomologischen Vereins. Supplement 8 (2001).

44. Ratnasingham, S., Hebert, P.D.N. BOLD: The Barcode of Life Data System (http://www.barcodinglife.org). Molecular Ecology Notes 7: 355–364 (2007). https://doi.org/10.1111/j.1471-8286.2006.01678.x

45. Kumar, S., Stecher, G., Li, M., Knyaz, C., Tamura, K. MEGA X: Molecular evolutionary genetics Analysis across computing platforms. Mol. Biol. Evol. 35, 1547–1549 (2018).

46. Nei, M., Tajima, F. DNA polymorphism detectable by restriction endonucleases. Genetics 97, 145–163 (1981).

47. Clement, M., Posada, D., Crandall, K. A. TCS: a computer program to estimate gene genealogies. Mol. Ecol. 9, 1657–1659 (2000).

48. ESRI. ArcGIS Desktop: Release 9.3. New York St., Redlands, CA. Environmental Systems Research Institute. http://www.esri.com/software/arcgis/eval-help/arcgis-93 (2008).

49. Kirichenko, N. The Lime Leafminer Phyllonorycter issikii in Western Siberia: some ecological characteristics of the population of the recent invader. Contemporary Problems of Ecology 7, 114–121 (2014).

