## Supplementary material for "Tracing the invasion of a leaf-mining moth in the Palearctic through DNA barcoding of historical herbaria"

**Table S1.** Findings of *Phyllonorycter messaniella* mines in herbarium specimens of *Tilia* spp. in Western Europe, 1915–1942*.

| № | Country | Region (city) | Number of | | | | | Year | *Tilia* species |
|---|---|---|---|---|---|---|---|---|---|
| | | | herba rium speci mens | mines | leaves with mines | leaves in herba-rium specimen | mined leaves per herbarium specimen, % | | |
| 1 | England | Hampshire | 1 | 1 | 1 | 10 | 10 | 1915 | *T. petiolaris* |
| 2 | England | Gloucester-shire | 1 | 3 | 3 | 12 | 25 | 1927 | *T. tomentosa* |
| 3 | England | Gloucester-shire | 1 | 1 | 1 | 8 | 13 | 1927 | *T. tomentosa* |
| 4 | Italy | Campania (Portichi) | 1 | 3 | 1 | 2 | 50 | 1927 | *T.* × *vulgaris*** |
| 5 | France | Normandy (Caen) | 1 | 1 | 1 | 2 | 50 | 1942 | *T. tomentosa* |
| 6 | France | Normandy (Caen, Venoix) | 1 | 2 | 1 | 1 | 100 | 1942 | *T. tomentosa* |
| | | In total: | 6 | 11 | 8 | | | | |

*The species identified by mines morphology, where possible, by pupal morphology and by DNA barcoding of larvae and pupae dissected from the mines. All herbarium specimens with *Ph. messaniella* mines are stored in Natural History Museum (London). ***T.* × *vulgaris* is presently considered a junior synonym of *T.* × *europaea* (which is a hybrid of *T. cordata* and *T. platyphyllos*).

**Table S2.** Findings of *Phyllonorycter lucetiella* and *Ph. tiliacella* mines in the herbarium specimens of *Tilia* spp. from North America, 1824–2010.

| № | Country | State | Number of | | | | | Year | Lime species | Deposi-taria |
|---|---|---|---|---|---|---|---|---|---|---|
| | | | herb. specimens | mines | leaves with mines | leaves in herbarium specimen | Mined leaves per herb. specimen, % | | | |
| ***Phyllonorycter lucetiella*** | | | | | | | | | | |
| 1 | Canada | Ontario | 1 | 1 | 1 | 5 | 20 | 1983 | *Tilia americana* | LECB |
| 2 | Canada | Ontario | 1 | 1 | 1 | 9 | 11 | 1962 | *T. americana* | W |
| 3 | Canada | Quebec | 1 | 1 | 1 | 8 | 13 | 1941 | *Tilia glabra* | MHA |
| 4 | USA | Alabama | 1 | 1 | 1 | 5 | 20 | 2008 | *T. cardiniana* | E |
| 5 | USA | Arkansas | 1 | 1 | 1 | 8 | 13 | 1972 | *Tilia* sp. | MHA |
| 6 | USA | Virginia | 1 | 1 | 1 | 6 | 17 | 1994 | *T. cardiniana* | E |
| 7 | USA | Illinois | 1 | 8 | 8 | 8 | 100 | 1979 | *T. americana* | TK |
| 8 | USA | Mississippi | 1 | 3 | 3 | 14 | 21 | 2000 | *T. americana* | E |
| 9 | USA | Mississippi | 1 | 1 | 1 | 10 | 10 | 2000 | *T. americana* | E |
| 10 | USA | Mississippi | 1 | 4 | 4 | 15 | 27 | 2000 | *T. americana* | E |
| 11 | USA | Mississippi | 1 | 1 | 1 | 6 | 17 | 2000 | *T. americana* | E |
| 12 | USA | Missouri | 1 | 2 | 2 | 8 | 25 | 1850 | *Tilia* sp. | G |
| 13 | USA | Missouri | 1 | 3 | 3 | 12 | 25 | 1922 | *Tilia americana* | E |
| 14 | USA | Missouri | 1 | 5 | 5 | 5 | 100 | 1921 | *Tilia americana* | K |
| 15 | USA | Missouri | 1 | 1 | 1 | 3 | 33 | 1921 | *Tilia americana* | K |
| 16 | USA | Missouri | 1 | 1 | 1 | 7 | 14 | 1988 | *T. americana* | K |
| 17 | USA | Michigan | 1 | 1 | 1 | 11 | 9 | 1994 | *T. americana* | E |
| 18 | USA | New York | 1 | 1 | 1 | 6 | 17 | 1994 | *T. americana* | W |
| 19 | USA | New York | 1 | 1 | 1 | 10 | 10 | 1948 | *T. heterophylla* | B |
| 20 | USA | New York | 1 | 1 | 1 | 9 | 11 | 2010 | *T. americana* | W |
| 21 | USA | New York | 1 | 1 | 1 | 12 | 8 | 1980 | *Tilia* sp. | LECB |
| 22 | USA | New York | 1 | 2 | 2 | 13 | 15 | 1980 | *Tilia* sp. | MHA |
| 23 | USA | New York | 1 | 3 | 3 | 16 | 19 | 1899 | *Tilia americana* | W |
| 24 | USA | N.Carolina | 1 | 2 | 2 | 5 | 40 | 1994 | *T. cardiniana* | E |
| 25 | USA | N.Carolina | 1 | 4 | 2 | 11 | 18 | 1994 | *T. cardiniana* | E |
| 26 | USA | N.Carolina | 1 | 8 | 5 | 8 | 63 | 1994 | *T. cardiniana* | E |
| 27 | USA | N.Carolina | 1 | 1 | 1 | 3 | 33 | 1994 | *T. cardiniana* | E |
| 28 | USA | N.Carolina | 1 | 7 | 5 | 10 | 50 | 1994 | *T. cardiniana* | E |
| 29 | USA | N.Carolina | 1 | 2 | 2 | 5 | 40 | 1994 | *T. cardiniana* | E |
| 30 | USA | N.Carolina | 1 | 10 | 10 | 11 | 91 | 1994 | *T. cardiniana* | E |
| 31 | USA | N.Carolina | 1 | 1 | 1 | 3 | 33 | 1994 | *T. cardiniana* | E |
| 32 | USA | Texas | 1 | 1 | 1 | 5 | 20 | 1872 | *T. americana* | E |
| 33 | USA | Florida | 1 | 2 | 2 | 7 | 29 | 2000 | *T. cardiniana* | E |
| 34 | USA | Florida | 1 | 1 | 1 | 7 | 14 | 2000 | *T. cardiniana* | E |
| | | **Total** | **34** | **84** | **77** | | | | | |
| ***Phyllonorycter tilicella*** | | | | | | | | | | |
| 35 | USA | Ohaio | 1 | 2 | 2 | 3 | 67 | 1905 | *Tilia* sp. | MHA |
| 36 | USA | Ohaio | 1 | 1 | 1 | 5 | 20 | 1924 | *T. heterophylla* | BM |
| 37 | USA | Penn. | 1 | 1 | 1 | 7 | 14 | 1824 | *T. americana* | W |
| | | **Total** | **3** | **4** | **4** | | | | | |

**Table S3.** Presence of *Phyllonorycter issikii* mines in the herbarium specimens of *Tilia* spp. collected in the last 253 years in the Palearctic.

| № | Lime species | Number of herbarium specimens with mines | | Number of mines | | Number of leaves with mines | Number of leaves in herbarium specimen | Number of mined leaves per herb. specimen, % |
|---|---|---|---|---|---|---|---|---|
| | | absolute value | % from all number of herbarium specimens with mines | absolute value | % from all mines | | | |
| **I. Putative invasive range (Europe, European part of Russia, Western Siberia)** | | | | | | | | |
| 1 | *T. cordata* | 15 | 7 | 102 | 7,9 | 62 | 300 | 21 |
| 2 | *T. platyphyllos* | 7 | 3 | 32 | 2,5 | 20 | 57 | 35 |
| **II. Putative native range (Russian Far East, Korea, China, Japan)** | | | | | | | | |
| 3 | *T. amurensis* | 74 | 33 | 661 | 51,3 | 327 | 1647 | 20 |
| 4 | *T. taquetii* | 42 | 19 | 312 | 24,2 | 197 | 1016 | 19 |
| 5 | *T. mandshurica* | 24 | 11 | 36 | 2,8 | 34 | 171 | 20 |
| 6 | *T. japonica* | 11 | 5 | 15 | 1,2 | 14 | 235 | 6 |
| 7 | *T. chinensis* | 8 | 4 | 11 | 0,9 | 11 | 135 | 8 |
| 8 | *T. maximowicziana* | 7 | 3 | 13 | 1,0 | 8 | 91 | 9 |
| 9 | *T. tuan* | 5 | 2 | 10 | 0,8 | 8 | 118 | 7 |
| 10 | *T. mongolica* | 5 | 2 | 5 | 0,4 | 5 | 101 | 5 |
| 11 | *T. laetevirens* | 3 | 1 | 26 | 2,0 | 20 | 28 | 71 |
| 12 | *T. kiusiana* | 2 | 1 | 5 | 0,4 | 5 | 31 | 16 |
| 13 | *T. pekinensis* | 1 | 0 | 3 | 0,2 | 2 | 6 | 33 |
| 14 | *T. intonsa* | 1 | 0 | 2 | 0,2 | 2 | 9 | 22 |
| 15 | *T. leptocarya* | 1 | 0 | 1 | 0,1 | 1 | 7 | 14 |
| 16 | *T. rufa* | 1 | 0 | 3 | 0,2 | 3 | 25 | 12 |
| 17 | *T. miqueliana* | 1 | 0 | 1 | 0,1 | 1 | 12 | 8 |
| 18 | *T. koreana* | 1 | 0 | 1 | 0,1 | 1 | 15 | 7 |
| 19 | *T. paucicostata* | 1 | 0 | 1 | 0,1 | 1 | 19 | 5 |
| 20 | *T. diviticata* | 1 | 0 | 1 | 0,1 | 1 | 25 | 4 |
| 21 | *T. sp* | 15 | 7 | 47 | 3,6 | 34 | 338 | 10 |
| | **TOTAL** | **226** | | **1288** | | **757** | **4386** | |

**Table S4.** Specimen data of DNA barcoded archival larvae and pupae of *Tilia*-feeding *Phyllonorycter* dissected from the mines in herbaria (dated by 1859–2014) in the Northern Hemisphere. The dataset is deposited in BOLD, doi (pending).

| № | *Phyllonorycter* species* | Stage** | Process ID | Collector (from herbarium) | Host plant*** | Country | Locality**** | Date of collection | Latitude | Longitude | Elevation, m |
|---|---|---|---|---|---|---|---|---|---|---|---|
| Archival *Phyllonorycter* specimens | | | | | | | | | | | |
| 1 | *Ph. issikii* | L | LMINH143-19 | N. Kirichenko | *T. cordata* | Germany | Hessen | 16.VII.2014 | 50.622 | 9.088 | 381 |
| 2 | *Ph. issikii* | L | LMINH144-19 | N. Kirichenko | *T. cordata* | Germany | Hessen | 16.VII.2014 | 50.622 | 9.088 | 381 |
| 3 | *Ph. issikii* | L | LMINH052-19 | N. Kirichenko | *T. platyphyllos* | Italy | Veneto | 04.X.2011 | 46.004 | 12.000 | 495 |
| 4 | *Ph. issikii* | P | LMINH020-19 | N. Kirichenko | *T. amurensis* | China | Jilin | 31.VIII.1951 | 43.111 | 126.301 | 396 |
| 5 | *Ph. issikii* | P | LMINH022-19 | N. Kirichenko | *T. mandshurica* | China | Shandong | 06.VIII.1924 | 35.665 | 117.710 | 201 |
| 6 | *Ph. issikii* | L | LMINH023-19 | N. Kirichenko | *T. taquetii* | China | Heilongjiang | 20.VI.1903 | 46.021 | 127.567 | 188 |
| 7 | *Ph. issikii* | L | LMINH026-19 | N. Kirichenko | *Tilia. sp.* | China | Heilongjiang | 26.VI.1903 | 46.146 | 129.178 | 439 |
| 8 | *Ph. issikii* | L | LMINH030-19 | N. Kirichenko | *T. amurensis* | China | Jilin | 29.VI.1896 | 44.159 | 123.852 | 167 |
| 9 | *Ph. issikii* | L | LMINH031-19 | N. Kirichenko | *T. amurensis* | China | Jilin | 29.VI.1896 | 44.159 | 123.852 | 167 |
| 10 | *Ph. issikii* | L | LMINH032-19 | N. Kirichenko | *T. amurensis* | China | Jilin | 29.VI.1896 | 44.159 | 123.852 | 167 |
| 11 | *Ph. issikii* | L | LMINH036-19 | N. Kirichenko | *T. laetevirens* | China | Gansu | 01.VI.1911 | 35.829 | 103.811 | 2435 |
| 12 | *Ph. issikii* | L | LMINH038-19 | N. Kirichenko | *T. mandshurica* | China | Jilin | 01.VII.1896 | 43.055 | 127.392 | 543 |
| 13 | *Ph. issikii* | P | LMINH001-19 | N. Kirichenko | *T. cordata* | Russia | KO | 19.VIII.2014 | 57.906 | 41.253 | 153 |
| 14 | *Ph. issikii* | L | LMINH002-19 | N. Kirichenko | *T. cordata* | Russia | KO | 27.VIII.2014 | 58.294 | 42.402 | 146 |
| 15 | *Ph. issikii* | P | LMINH003-19 | N. Kirichenko | *T. cordata* | Russia | ChO | 28.VII.1987 | 55.159 | 59.615 | 601 |
| 16 | *Ph. issikii* | L | LMINH004-19 | N. Kirichenko | *T. cordata* | Russia | SO | 29.VIII.1990 | 53.433 | 49.672 | 98 |
| 17 | *Ph. issikii* | L | LMINH006-19 | N. Kirichenko | *T. amurensis* | Russia | PK | 17.VII.1951 | 43.579 | 131.992 | 63 |
| 18 | *Ph. issikii* | P | LMINH007-19 | N. Kirichenko | *T. amurensis* | Russia | PK | 17.VII.1951 | 43.579 | 131.992 | 63 |
| 19 | *Ph. issikii* | P | LMINH008-19 | N. Kirichenko | *T. amurensis* | Russia | PK | 17.VII.1951 | 43.579 | 131.992 | 63 |
| 20 | *Ph. issikii* | L | LMINH009-19 | N. Kirichenko | *T. amurensis* | Russia | PK | 30.VII.1952 | 43.315 | 132.517 | 237 |
| 21 | *Ph. issikii* | L | LMINH010-19 | N. Kirichenko | *T. amurensis* | Russia | PK | 25.VII.1952 | 43.654 | 132.524 | 211 |
| 22 | *Ph. issikii* | L | LMINH011-19 | N. Kirichenko | *T. amurensis* | Russia | PK | 26.VII.1950 | 43.669 | 132.527 | 452 |
| 23 | *Ph. issikii* | L | LMINH012-19 | N. Kirichenko | *T. amurensis* | Russia | AO | 02.VIII.1914 | 54.636 | 126.748 | 509 |
| 24 | *Ph. issikii* | L | LMINH013-19 | N. Kirichenko | *T. amurensis* | Russia | PK | 24.VII.1936 | 43.699 | 132.169 | 236 |
| 25 | *Ph. issikii* | L | LMINH014-19 | N. Kirichenko | *T. taquetii* | Russia | PK | 14.VII.1951 | 43.696 | 132.167 | 211 |

| № | *Phyllonorycter* species* | Stage** | Process ID | Collector (from herbarium) | Host plant*** | Country | Locality**** | Date of collection | Latitude | Longitude | Elevation, m |
|---|---|---|---|---|---|---|---|---|---|---|---|
| 26 | *Ph. issikii* | L | LMINH015-19 | N. Kirichenko | *T. taquetii* | Russia | PK | 14.VII.1951 | 43.696 | 132.167 | 211 |
| 27 | *Ph. issikii* | L | LMINH016-19 | N. Kirichenko | *T. taquetii* | Russia | PK | 17.VII.1936 | 43.669 | 132.527 | 452 |
| 28 | *Ph. issikii* | L | LMINH017-19 | N. Kirichenko | *T. taquetii* | Russia | PK | 21.VII.1936 | 43.669 | 132.527 | 452 |
| 29 | *Ph. issikii* | P | LMINH018-19 | N. Kirichenko | *T. taquetii* | Russia | PK | 28.VII.1936 | 43.669 | 132.527 | 452 |
| 30 | *Ph. issikii* | P | LMINH019-19 | N. Kirichenko | *T. taquetii* | Russia | PK | 28.VII.1936 | 43.669 | 132.527 | 452 |
| 31 | *Ph. issikii* | L | LMINH041-19 | N. Kirichenko | *T. taquetii* | Russia | PK | 01.VI.1981 | 43.854 | 135.152 | 179 |
| 32 | *Ph. issikii* | L | LMINH042-19 | N. Kirichenko | *T. taquetii* | Russia | PK | 01.VI.1936 | 43.669 | 132.527 | 452 |
| 33 | *Ph. issikii* | P | LMINH046-19 | N. Kirichenko | *T. amurensis* | Russia | PK | 01.VI.1936 | 43.669 | 132.527 | 452 |
| 34 | *Ph. issikii* | L | LMINH047-19 | N. Kirichenko | *T. amurensis* | Russia | PK | 01.VI.1941 | 45.734 | 135.156 | 439 |
| 35 | *Ph. issikii* | L | LMINH055-19 | N. Kirichenko | *T. taquetii* | Russia | PK | 01.VII.1951 | 43.579 | 131.992 | 63 |
| 36 | *Ph. issikii* | L | LMINH098-19 | N. Kirichenko | *T. mandshurica* | Russia | PK | 16.VII.1968 | 43.316 | 132.684 | 267 |
| 37 | *Ph. issikii* | L | LMINH103-19 | N. Kirichenko | *T. amurensis* | Russia | PK | 24.VI.1987 | 43.177 | 131.970 | 184 |
| 38 | *Ph. issikii* | L | LMINH104-19 | N. Kirichenko | *T. amurensis* | Russia | PK | 04.VII.1998 | 43.692 | 132.155 | 147 |
| 39 | *Ph. issikii* | L | LMINH105-19 | N. Kirichenko | *T. amurensis* | Russia | PK | 17.VII.1992 | 43.197 | 132.111 | 8 |
| 40 | *Ph. issikii* | P | LMINH106-19 | N. Kirichenko | *T. amurensis* | Russia | PK | 17.VII.1992 | 43.197 | 132.111 | 8 |
| 41 | *Ph. issikii* | P | LMINH107-19 | N. Kirichenko | *T. amurensis* | Russia | KhK | 25.VIII.1985 | 50.345 | 137.719 | 92 |
| 42 | *Ph. issikii* | L | LMINH108-19 | N. Kirichenko | *T. amurensis* | Russia | KhK | 01.VI.1961 | 49.991 | 135.304 | 269 |
| 43 | *Ph. issikii* | L | LMINH109-19 | N. Kirichenko | *T. amurensis* | Russia | KhK | 01.VI.1961 | 49.991 | 135.304 | 269 |
| 44 | *Ph. issikii* | L | LMINH110-19 | N. Kirichenko | *T. amurensis* | Russia | KhK | 01.VI.1961 | 49.991 | 135.304 | 269 |
| 45 | *Ph. issikii* | L | LMINH111-19 | N. Kirichenko | *T. amurensis* | Russia | KhK | 06.VIII.1977 | 54.164 | 126.157 | 851 |
| 46 | *Ph. issikii* | P | LMINH113-19 | N. Kirichenko | *T. heterophylla* | Russia | PK | 23.VIII.1975 | 42.483 | 130.750 | 3 |
| 47 | *Ph. issikii* | P | LMINH119-19 | N. Kirichenko | *T. heterophylla* | Russia | AO | 23.VII.1859 | 54.021 | 126.743 | 686 |
| 48 | *Ph. issikii* | L | LMINH147-19 | N. Kirichenko | *T. taquetii* | Russia | PK | 18.VII.1951 | 43.634 | 132.488 | 343 |
| 49 | *Ph. issikii* | L | LMINH039-19 | N. Kirichenko | *T. maximowicziana* | Japan | Hokkaido | 01.VI.1905 | 43.349 | 141.639 | 171 |
| 50 | *Ph. issikii* | L | LMINH123-19 | N. Kirichenko | *T. maximowicziana* | Japan | Hokkaido | 19.VII.1956 | 42.927 | 141.125 | 598 |
| 51 | *Ph. messaniella* | L | LMINH121-19 | N. Kirichenko | *T. vulgaris* | Italy | Campania, Portichi | 12.V.1927 | 40.812 | 14.345 | 59 |
| 52 | *Ph. messaniella* | P | LMINH122-19 | N. Kirichenko | *T. tomentosa* | France | Normandia, Caen | 12.IX.1942 | 49.163 | -0.370 | 10 |
| 53 | *Ph. tiliacella* | P | LMINH120-19 | N. Kirichenko | *T. heterophylla* | USA | Ohaio | 01.VI.1960 | 40.616 | -82.879 | 320 |

| № | *Phyllonorycter* species* | Stage** | Process ID | Collector (from herbarium) | Host plant*** | Country | Locality**** | Date of collection | Latitude | Longitude | Elevation, m |
|---|---|---|---|---|---|---|---|---|---|---|---|
| 54 | *Ph. tiliacella* | P | LMINH153-19 | N. Kirichenko | *T. americana* | USA | Pennsylvania | 01.VIII.1894 | 39.843 | -77.960 | 334 |
| 55 | *Ph. lucetiella* | L | LMINH040-19 | N. Kirichenko | *T. glabra* | Canada | Ontario | 01.VI.1941 | 45.444 | -75.611 | 84 |
| 56 | *Ph. lucetiella* | L | LMINH043-19 | N. Kirichenko | *T. americana* | USA | Massachusetts | 17.IX.1979 | 42.499 | -71.546 | 42 |
| 57 | *Ph. lucetiella* | L | LMINH044-19 | N. Kirichenko | *T. americana* | USA | Massachusetts | 17.IX.1979 | 42.499 | -71.546 | 42 |
| 58 | *Ph. lucetiella* | L | LMINH045-19 | N. Kirichenko | *T. americana* | USA | Massachusetts | 17.IX.1979 | 42.499 | -71.546 | 42 |
| 59 | *Ph. lucetiella* | L | LMINH145-19 | N. Kirichenko | *T. heterophylla* | USA | New York | 15.VII.1948 | 42.742 | -76.597 | 309 |
| 60 | *Ph. lucetiella* | L | LMINH152-19 | N. Kirichenko | *T. americana* | USA | New York | 01.VI.2010 | 40.731 | -73.991 | 5 |
| 61 | *Ph. lucetiella* | L | LMINH128-19 | N. Kirichenko | *T. caroliniana* | USA | N.Carolina | 08.X.1994 | 34.540 | -77.397 | 1 |
| 62 | *Ph. lucetiella* | L | LMINH129-19 | N. Kirichenko | *T. caroliniana* | USA | N.Carolina | 08.X.1994 | 34.540 | -77.397 | 1 |
| 63 | *Ph. lucetiella* | L | LMINH130-19 | N. Kirichenko | *T. caroliniana* | USA | N.Carolina | 08.X.1994 | 34.540 | -77.397 | 1 |
| 64 | *Ph. lucetiella* | L | LMINH131-19 | N. Kirichenko | *T. caroliniana* | USA | N.Carolina | 08.X.1994 | 34.540 | -77.397 | 1 |
| 65 | *Ph. lucetiella* | L | LMINH132-19 | N. Kirichenko | *T. caroliniana* | USA | N.Carolina | 08.X.1994 | 34.540 | -77.397 | 1 |
| 66 | *Ph. lucetiella* | L | LMINH133-19 | N. Kirichenko | *T. caroliniana* | USA | Florida | 11.VIII.2000 | 30.521 | -84.760 | 38 |
| 67 | *Ph. lucetiella* | P | LMINH137-19 | N. Kirichenko | *T. caroliniana* | USA | Florida | 07.VIII.2000 | 30.575 | -85.085 | 36 |
| 68 | *Phyllonorycter* sp. | L | LMINH096-19 | N. Kirichenko | *T. amurensis* | Russia | PK | 01.VIII.1995 | 42.765 | 132.335 | 256 |
| 69 | *Phyllonorycter* sp. | L | LMINH099-19 | N. Kirichenko | *T. taquetii* | Russia | PK | 03.X.1997 | 43.006 | 131.848 | 127 |
| 70 | *Phyllonorycter* sp. | P | LMINH101-19 | N. Kirichenko | *T. taquetii* | Russia | PK | 03.X.1997 | 43.006 | 131.848 | 127 |
| 71 | *Phyllonorycter* sp. | L | LMINH102-19 | N. Kirichenko | *T. taquetii* | Russia | PK | 24.VI.1987 | 43.417 | 133.485 | 752 |
| | | | | | **Reference DNA barcodes** | | | | | | |
| 1 | *Ph. issikii* | I | PHLAD034-11 | P. Huemer | — | Италия | South Tyrol | 14.IX.2011 | 46.428 | 11.300 | 643 |
| 2 | *Ph. issikii* | I | ISSIK001-12 | N. Kirichenko | *T. cordata* | Russia | MO | 21.VI.2010 | 54.839 | 37.604 | 162 |
| 3 | *Ph. issikii* | L | ISSIK269-14 | N. Kirichenko | *T. taquetii* | Russia | PK | 28.VIII.2011 | 43.689 | 132.157 | 160 |
| 4 | *Ph. issikii* | L | ISSIK378-15 | N. Kirichenko | *T. mandshurica* | Russia | PK | 17.VII.2013 | 43.681 | 132.160 | 224 |
| 5 | *Ph. issikii* | I | GRAAM048-13 | G. Deschka | *T. maximowicziana* | Japan | Hokkaido | 28.IX.1966 | 43.113 | 140.453 | 30 |
| 6 | *Ph. issikii* | I | ISSIK297-14 | K. Tokashi | *T. maximowicziana* | Japan | Hokkaido | 13.IX.2014 | 43.034 | 141.315 | 121 |
| 7 | *Ph. issikii* | I | ISSIK317-14 | K. Tokashi | *T. maximowicziana* | Japan | Hokkaido | 13.IX.2014 | 43.034 | 141.315 | 121 |
| 8 | *Ph. messaniella* | I | GRPAL161-11 | A. Cama | — | France | Loire | 12.XII\|.2006 | 47.250 | 0.210 | 33 |
| 9 | *Ph. lucetiella* | P | MICRU060-15 | J.-F. Landry | *T. americana* | Canada | Quebec | 27.IX.2015 | 45.469 | -75.811 | 180 |

| № | ***Phyllonorycter* species**[*] | **Stage**[**] | **Process ID** | **Collector (from herbarium)** | **Host plant**[***] | **Country** | **Locality**[****] | **Date of collection** | **Latitude** | **Longitude** | **Elevation, m** |
|---|---|---|---|---|---|---|---|---|---|---|---|
| 10 | *Phyllonorycter* sp. | L | ISSIK371-14 | K. Tokashi | *T. japonica* | Japan | Honshu | 01.IX.2014 | 38.260 | 140.850 | 244 |
| Outgroup | | | | | | | | | | | |
| 11 | *Tischeria* sp. | L | ISSIK102-14 | N. Kirichenko | *T. taquetii* | Russia | PK | 25.VIII.2011 | 43.689 | 132.157 | 160 |

[*]Species: sp. – putative new species; [**]Stage: L – larva, P – pupa, I – imago; [***]Host plant: — no data available; [****]Localities: SO – Sverdlovsk Oblast, KO – Kostroma Oblast, ChO – Chelyabinsk Oblast, AO – Amur Oblast, PK – Primorskiy Krai, KhK – Khabarovsk Krai.

**Table S5.** The historical haplotypes of *Phyllonorycter issikii* (COI gene mtDNA) obtained in the present study and their correspondence to those in the modern range of the species in the Palearctic[20].

| Haplo-type number | Country, region* | Years | Obtained sequence length, bp. | Process ID | Presence of the haplotype in the modern *Ph. issikii* range**,20 |
|---|---|---|---|---|---|
| H1 | Russia, RFE | 1951 | 563 | LMINH007-19 | Austria, Bulgaria, Hungary, Germany, Netherlands, Poland, Russia (European part), Slovenia, Ukraine, Finland, Czech Republic, Japan (Hokkaido) |
| | Russia, RFE | 1951 | 563 | LMINH006-19 | |
| | Russia, RFE | 1952 | 563 | LMINH009-19 | |
| | Russia, RFE | 1950 | 563 | LMINH011-19 | |
| | Russia, RFE | 1936 | 563 | LMINH013-19 | |
| | Russia, RFE | 1951 | 563 | LMINH015-19 | |
| | Russia, RFE | 1936 | 563 | LMINH016-19 | |
| | Russia, RFE | 1936 | 511 | LMIMH019-19 | |
| | Russia, RFE | 1951 | 511 | LMIMH055-19 | |
| | Japan, Hokkaido | 1956 | 563 | LMINH123-19 | |
| | Russia, ER | 1987 | 563 | LMINH003-19 | |
| | Germany | 2014 | 563 | LMINH143-19 | |
| | Germany | 2014 | 563 | LMINH144-19 | |
| H2 | Russia, RFE | 1981 | 563 | LMINH041-19 | Russia (Siberia) |
| H8 | Russia, RFE | 1936 | 563 | LMINH018-19 | Austria, Bulgaria, Hungary, Germany, Italy, Lithuania, Russia (European part, Siberia, RFE), Slovenia, Ukraine, Finland |
| | Italy | 2011 | 563 | LMINH052-19 | |
| H13 | Russia, RFE | 1951 | 563 | LMINH008-19 | Russia (Siberia) |
| | Russia, RFE | 1992 | 563 | LMINH105-19 | |
| H22 | China, Heilongjiang | 1903 | 278 | LMINH023-19 | Russia (European part) |
| H23 | Russia, RFE | 1987 | 563 | LMINH103-19 | Austria, Bulgaria, Hungary, Germany, Lithuania, Netherlands, Poland, Russia (European part, Siberia), Ukraine, Finland |
| | Russia, RFE | 1951 | 563 | LMINH014-19 | |
| | Russia, RFE | 1992 | 563 | LMIMH106-19 | |
| | Russia, RFE | 1951 | 563 | LMINH147-19 | |
| | Russia, RFE | 1961 | 563 | LMINH108-19 | |
| | Russia, ER | 2014 | 563 | LMINH001-19 | |
| | Russia, ER | 1990 | 563 | LMINH004-19 | |
| H26 | China, Jilin | 1951 | 563 | LMINH020-19 | Russia (RFE) |
| H28 | Russia, RFE | 1975 | 513 | LMINH113-19 | Russia (RFE) |
| H30 | Russia, RFE | 1998 | 563 | LMINH104-19 | Russia (RFE) |

*RFE – Russian Far East, ER – European Russia; **Countries and regions where the same haplotypes have been found in the historical and modern areas are underlined.

**Table S6.** The distribution of old haplotypes of *Phyllonorycter issikii* (COI mtDNA) in different countries and regions across the Palearctic based on data from archival herbaria.

| Haplotype number | Country, region [1] | Years | Obtained sequence length, bp | Lime species | Process ID |
|---|---|---|---|---|---|
| 1 | Russia, RFE (PK) | 1951 | 563 | *T. maximowicziana* | LMINH007-19 |
| 2 | Russia, RFE (KhK) | 1985 | 563 | *T. amurensis* | LMINH107-19 |
| 3 | Russia, RFE (PK) | 1952 | 379 | *T. amurensis* | LMINH010-19 |
| 4 | Russia, RFE (KhK) | 1961 | 458 | *T. amurensis* | LMINH109-19 |
| 5 | Russia, RFE (KhK) | 1977 | 563 | *T. amurensis* | LMINH111-19 |
| 6 | Russia, RFE (AO) | 1859 | 408 | *T. amurensis* | LMINH119-19 |
| 7 | Russia, RFE (KhK) | 1961 | 563 | *T. amurensis* | LMINH110-19 |
| 8 | Japan, Hokkaido | 1905 | 133 | *T. maximowicziana* | LMINH039-19 |
| 9 | China, Jilin | 1896 | 133 | *T. mandshurica* | LMINH038-19 |
| | China, Heilongjiang | 1903 | 563 | *Tilia* sp. | LMINH026-19 |
| 10 | China, Jilin | 1896 | 278 | *T. amurensis* | LMINH030-19 |
| 11 | China, Julin | 1896 | 131 | *T. amurensis* | LMINH032-19 |
| 12 | Russia, RFE (PK) | 1941 | 563 | *T. amurensis* | LMINH047-19 |
| 13 | Russia, ER (KO) | 2014 | 563 | *T. cordata* | LMINH002-19 |
| 14 | Russia, RFE (PK) | 1936 | 563 | *T. taquetii* | LMINH017-19 |
| 15 | Russia, RFE (PK) | 1968 | 458 | *T. taquetii* | LMINH098-19 |
| 16 | China, Shandong | 1924 | 563 | *T. mandshurica* | LMINH022-19 |

[1]RFE – Russian Far East, PK – Primorsky Krai, KhK – Khabarovsk Krai, AO – Amur Oblast; ER – European Russia, KO – Kostroma Oblast. For China, provinces are indicated.

**Table S7.** The studied number of herbarium specimens of *Tilia* spp. from the Northern Hemisphere stored in 20 herbarium depositaria in Eurasia [*].

| № | Herbarium institution, city, country | Herba-rium code | Number of herbarium specimens from | |
|---|---|---|---|---|
| | | | Palearctic | Nearctic |
| 1 | National Museum of Natural History, Paris, France[**] | P | 1926 | 2 |
| 2 | Komarov Botanical Institute of the Russian Academy of Sciences, Saint Petersburg, Russia[**] | LECB | 2854 | 33 |
| 3 | Royal Botanic Gardens, Kew, London, UK | R | 1870 | 376 |
| 4 | Naturalis Biodiversity Center, Leiden, the Netherlands | L | 607 | 32 |
| 5 | Natural History Museum, London, UK | BM | 1628 | 104 |
| 6 | Conservatory and Botanical Garden of the city of Geneva, Geneva, Switzerland | G | 1625 | 5 |
| 7 | Natural History Museum, Vienna, Austria | W | 1357 | 74 |
| 8 | Berlin Botanic Garden and Botanical Museum, Berlin, Germany | B | 1134 | 26 |
| 9 | Botanical Garden Zürich, Zurich, Switzerland | ZT | 871 | 22 |
| 10 | Moscow State University, Moscow, Russia | MW | 761 | 22 |
| 11 | Natural History Museum, Florence, Italy | Fl | 671 | 6 |
| 12 | Royal Botanic Garden Edinburgh, Edinburgh, Scotland | E | 1674 | 237 |
| 13 | Principle Botanical Garden RAS, Moscow, Russia | MHA | 539 | 23 |
| 14 | Sapienza University of Rome, Rome, Italy | RO | 321 | 0 |
| 15 | National Museums Collection Center, Edinburgh, Scotland | — | 7 | 0 |
| 16 | Federal Scientific Center of the East Asia Terrestrial Biodiversity FEB RAS, Vladivostok, Russia | VLA | 444 | 0 |
| 17 | Botanic Garden–Institute FEE RAS (Far Eastern Branch of the Russian Academy of Sciences), Vladivostok, Russia | VBGI | 405 | 0 |
| 18 | Tomsk State University, Tomsk, Russia | TK | 253 | 6 |
| 19 | Central Siberian Botanical Garden SB RAS (Siberian Branch of the Russian Academy of Sciences), Novosibirsk, Russia | NS | 180 | 0 |
| 20 | Sukachev Institute of Forest SB RAS, Federal Research Center «Krasnoyarsk Science Center SB RAS», Krasnoyarsk, Russia | KRF | 8 | 0 |
| | TOTAL | | 19135 | 968 |

[*]The depositaria are listed in an approximate order according to overall herbarium size.[**]The biggest herbaria in the world.

**Table S8.** *Tilia* species studied in the herbarium collections from different biogeographic regions[*].

| **Lime species**[**] | **Number of lime species** |
|---|---|
| **Palearctic** | |
| *Tilia alba,* ***T. amurensis,*** *T. apiculata, T. argentea, T. asplenifolia, T. aurea, T. blockiana, T. budensis, T. calvescens, T. caucasica, T. chinensis, T. chingiana, T. concinna,* ***T. cordata,*** *T. cordifolia, T. dasystyla, T. divaricata, T. dictyoneura, T. endochrysea, T. euchlora, T. flava, T. flavescens, T. floribunda, T. furedensis, T. gizellae, T. glabra, T. grandifolia, T. haringiana, T. haszlinszkyana, T. haynaldiana, T. henryana, T. heterophylla, T. insularis, T. intercedens, T. intermedia, T. intonsa, T. japonica, T. kiusiana, T. komarovi, T. koreana, T. laetevirens, T. latebracteata, T. latifolia, T. laxiflora, T. ledebourii, T. leptocarya,* ***T. mandshurica, T. maximowicziana****, T. microphylla, T. miqueliana, T. mollis, T. moltkei, T. mongolica, T. mutabilis, T. nasczokinii, T. neglecta, T. nickerlii, T. nobilis, T. oblique, T. oblongifolia T. obricularis, T. oliveri, T. oxycarpa, T. pallida, T. parvifolia, T. paucicostata, T. pekinensis, T. perneckensis, T. pilosa,* ***T. platyphyllos,*** *T. praecox, T. pubescens, T. pyramidalis, T. rubescens, T. rubra, T. rufa, T. ruprechtii, T. sibirica T. septemtrionalis, T. spectabilis, T. sphaerocarpa, T. stenocarpa, T. stohlii, T. subangulata, T. subflavescens, T. sublanata, T. sylvestris, T. sythensis,* ***T. taquetii,*** *T. tennifolia, T. tomentosa, T. trichoclados, T. truncate, T. tuan, T. tucekii, T. turbinata, T. ulmifolia, T. vestita, T. viridis, T. vitifolia, T. vulgaris* | 101 |
| *Tilia* spp. (hybrids) | > 80 |
| **Nearctic** | |
| ***T. americana,*** *T. californiana, T. canadensis, T. caroliniana, T. floridana, T. glabra, T. heterophylla, T. lasioclada, T. leptophylla, T. littoralis, T. mexicana, T. michauxii, T. monticola, T. nuda, T. relicta, T. venulosa* | 16 |
| **Total number of species (excluding hybrids)** | 117[***] |

[*]Based on examination of herbarium specimens in 20 herbarium depositaria (see Table S7).

[**]The taxonomy of limes has been revised several times: *Tilia divaricata, T. koreana* and T. *taquetii* are presently considered as synonyms of *T. amurensis*; *T. sibirica* as a synonym of *T. cordata*; *T. pekinensis* as a synonym of *T. mandshurica*[42]. In out study, the names of lime species are provided as they were indicated on the labels at herbarium specimens. *Tilia* species marked in bold were dominant in herbaria (altogether they accounted 50% of all studied herbarium specimens).

**Table S9.** Characteristics of leaf mines and pupal cremaster in the Palearctic and Nearctic *Tilia*-feeding *Phyllonorycter* species*

| Species | Mine characteristics | | | Pupal cremaster |
|---|---|---|---|---|
| | **shape** | **position** | **folds on the epidermis covering mine** | |
| **The Palearctic species** | | | | |
| *Ph. issikii* | oval blotch | lower side of the leaf usually between two secondary veins | without folds | roundish, with one pair of spines having wide base and curved outward tips |
| *Ph. messaniella* | oval blotch | lower side of the leaf, usually between two veins | one distinct longitude fold which remains well visible in pressed leaves | square-like, with two pairs of spines: one pair are long spines with relatively narrow base, another pair are short straight spine |
| **The Nearctic species** | | | | |
| *Ph. lucetiella* | somewhat triangular blotch | lower side of the eaf in the angle between the two veins | without folds | trapezoid, with two pairs of small spines: one pair curved outwards and another pair curved inward |
| *Ph. tiliacella* | irregular blotch | usually upper side of the leaf | without folds but with circular lines or dots of frass on the epidermis covering mine | unknown |

*According to published data[14,31,32,43]

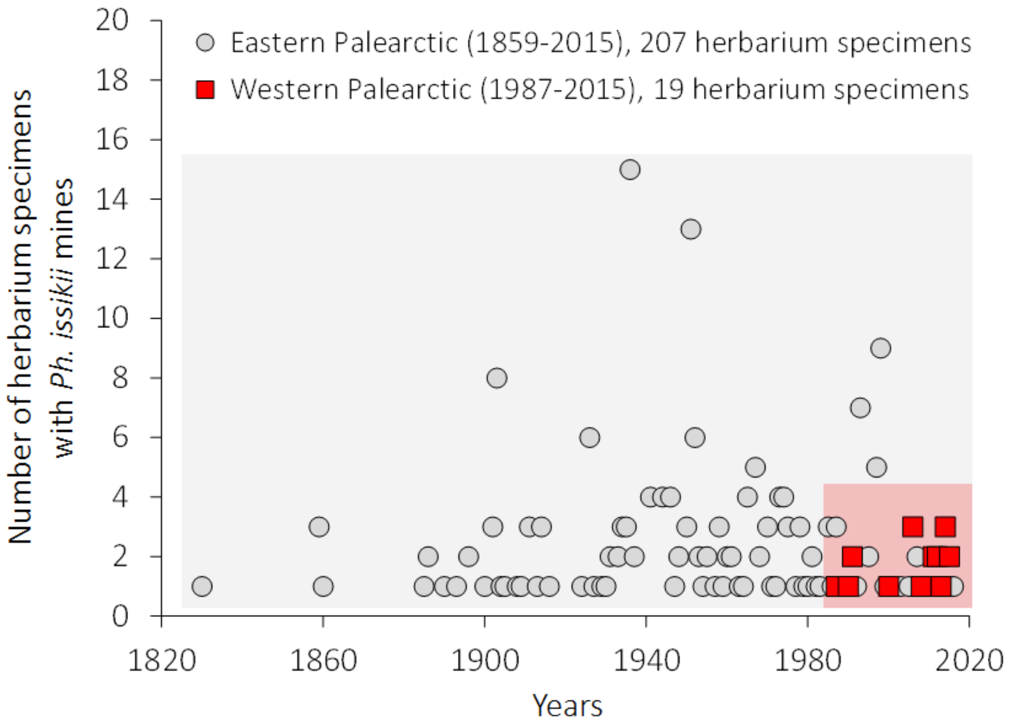


**Figure S1.** Dating of *Phyllonorycter issikii* mines in the herbariums specimens in the Palearctic: on the East, the putative native range (gray block) and on the West, the putative invasive range (red block).

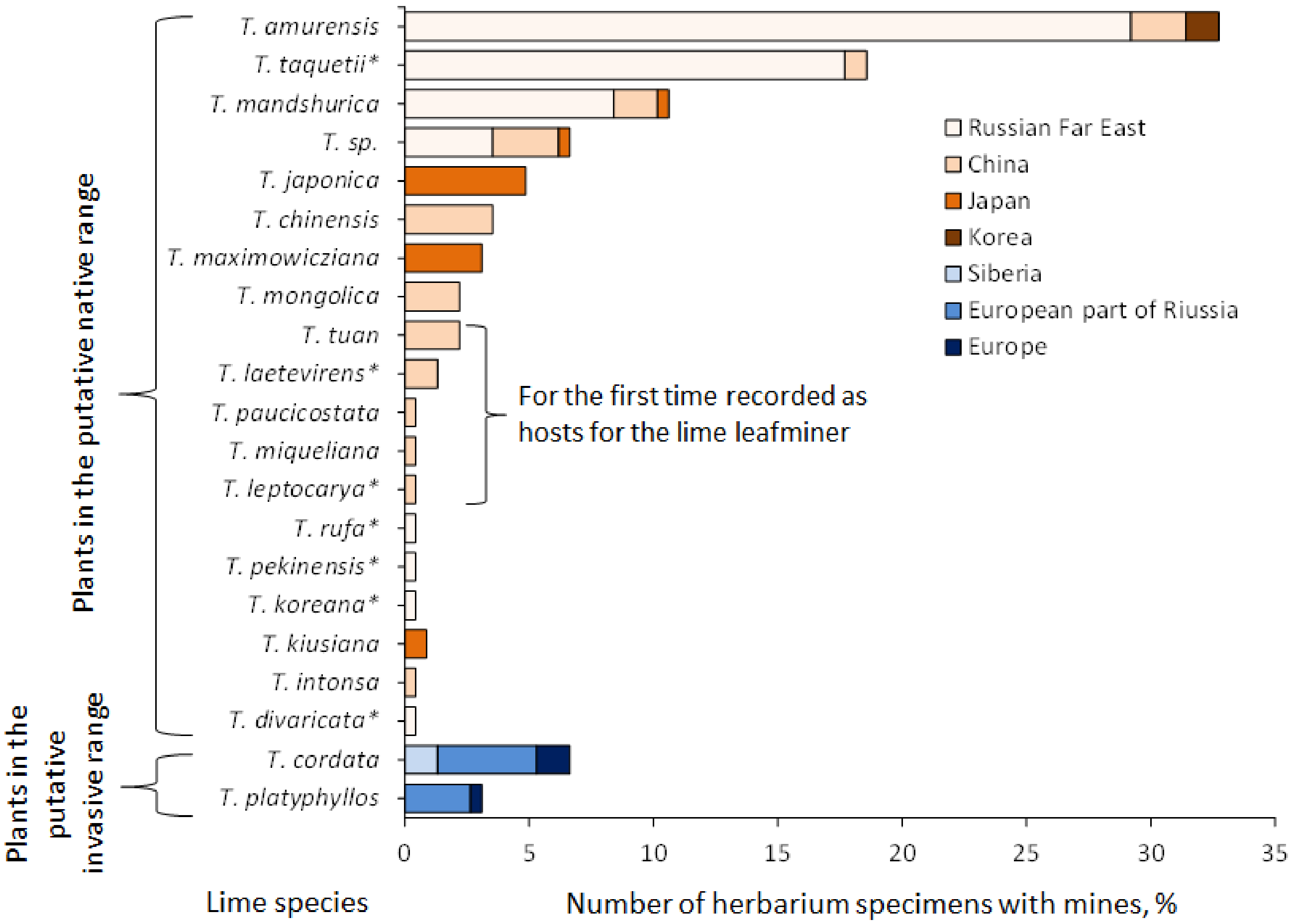


**Figure S2.** Trophic associations of *Phyllonorycter issikii* with different *Tilia* spp. based on findings of mines in historical herbaria collected in East Asia (the putative native range) and Western Palearctic (invasive range) over the past 253 years. The species marked by asterisk are presently known as junior synonyms: *Tilia taquetii, T. divaricata, T. koreana, T. rufa* are junior synonyms of *T. amurensis*; *T. laetevirens* is such of *T. chinensis*; *T. leptocarya* is such of *T. endochrysea*; *T. pekinensis* is such of *T. mandshurica.*

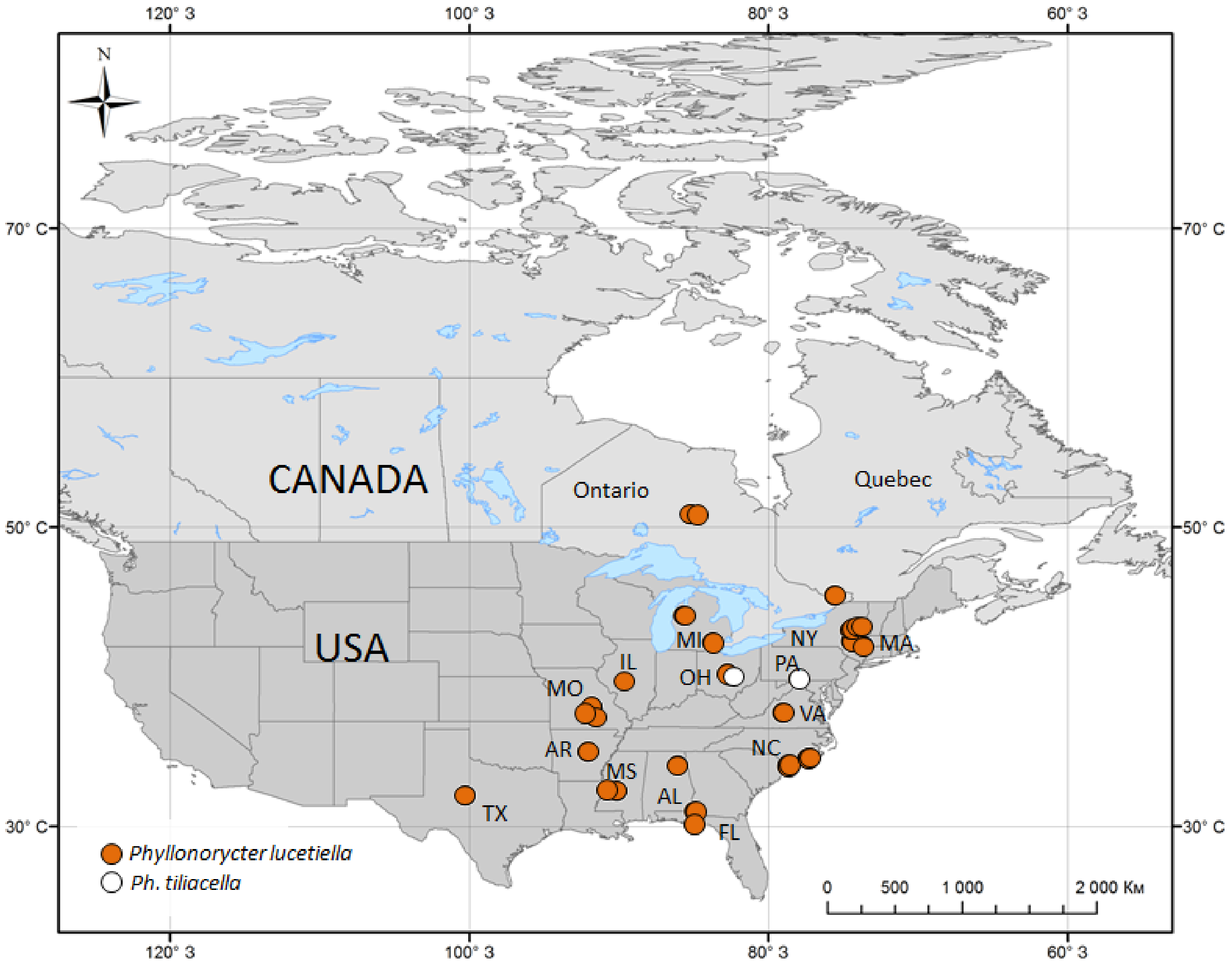


**Figures S3.** The findings of *Phyllonorycter lucetiella* and *Ph. tiliacella* mines in the herbarium specimens of *Tilia* collected in North America in the last 200 years. The states: AL – Alabama, AR – Arkansas, FL – Florida, IL – Illinois, MI – Michigan, MA – Massachusetts, MS – Mississippi, MO – Missouri, NC – North Carolina, NY – New York, OH – Ohio, PA – Pennsylvania, TX – Texas, VA – Virginia.

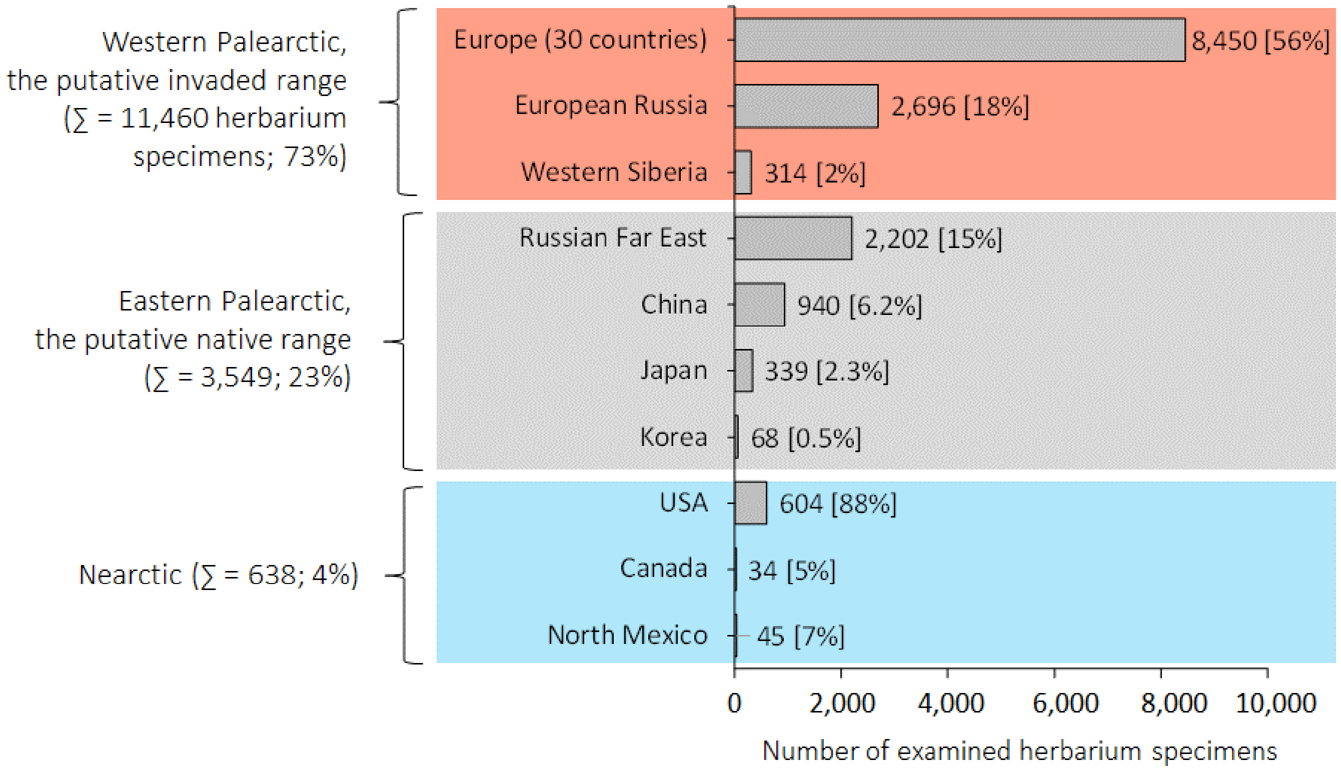


**Figure S4.** The number of herbarium specimens of *Tilia* from the Northern Hemisphere involved in the study. The red block indicates the part of the Palearctic where *Phyllonorycter issikii* is considered an invasive species, gray block where it is known as native species. The number of studied herbarium specimens are given in round brackets on the left side of the graph. The number of herbarium specimens and their percentage (%) in a particular region or a country is given in square brackets next to corresponding column. A herbarium specimen accounted from 1 to 72 lime leaves.
